# Bioengineered 3D hPSC-cholangiocyte ducts with physiological signals for biliary disease modelling

**DOI:** 10.1101/2025.11.10.686855

**Authors:** Britney Tian, Mina Ogawa, Manami Kondo, Gabriella Langeveld, Ling Jun Huan, Feng Zhang, Andrew Hollinger, Sara Deir, Christine Bear, Boyang Zhang, Shinichiro Ogawa

## Abstract

The progression of intrahepatic biliary diseases remains poorly understood, underscoring the urgent need to develop physiologically relevant human intrahepatic cholangiocyte disease models. Current approaches lack the complexity and throughput to capture how diverse biliary microenvironmental signals shape cholangiocyte behavior. To address this gap, we created a fully epithelialized and perfusible 3D bile duct from human pluripotent stem cell-derived cholangiocytes characterized by robust primary ciliation, apical-basal polarity and CFTR-mediated chloride conductance. Our results revealed physiologically relevant fluid flow and biliary stroma cells as essential components for sustaining cholangiocyte epithelial barrier integrity and ciliation. From here, we interrogated a broad spectrum of biliary signals to study bile acid toxicity and cytokine-driven injury, offering an unprecedented view into intrahepatic cholangiocyte stress responses and potential pathological mechanisms. By integrating physiologically relevant signals, fidelity and functional resolution, this platform provides the foundation needed to precisely decode pathogenic drivers and accelerate therapeutic development for devastating cholangiopathies.

**Graphical Abstract:** 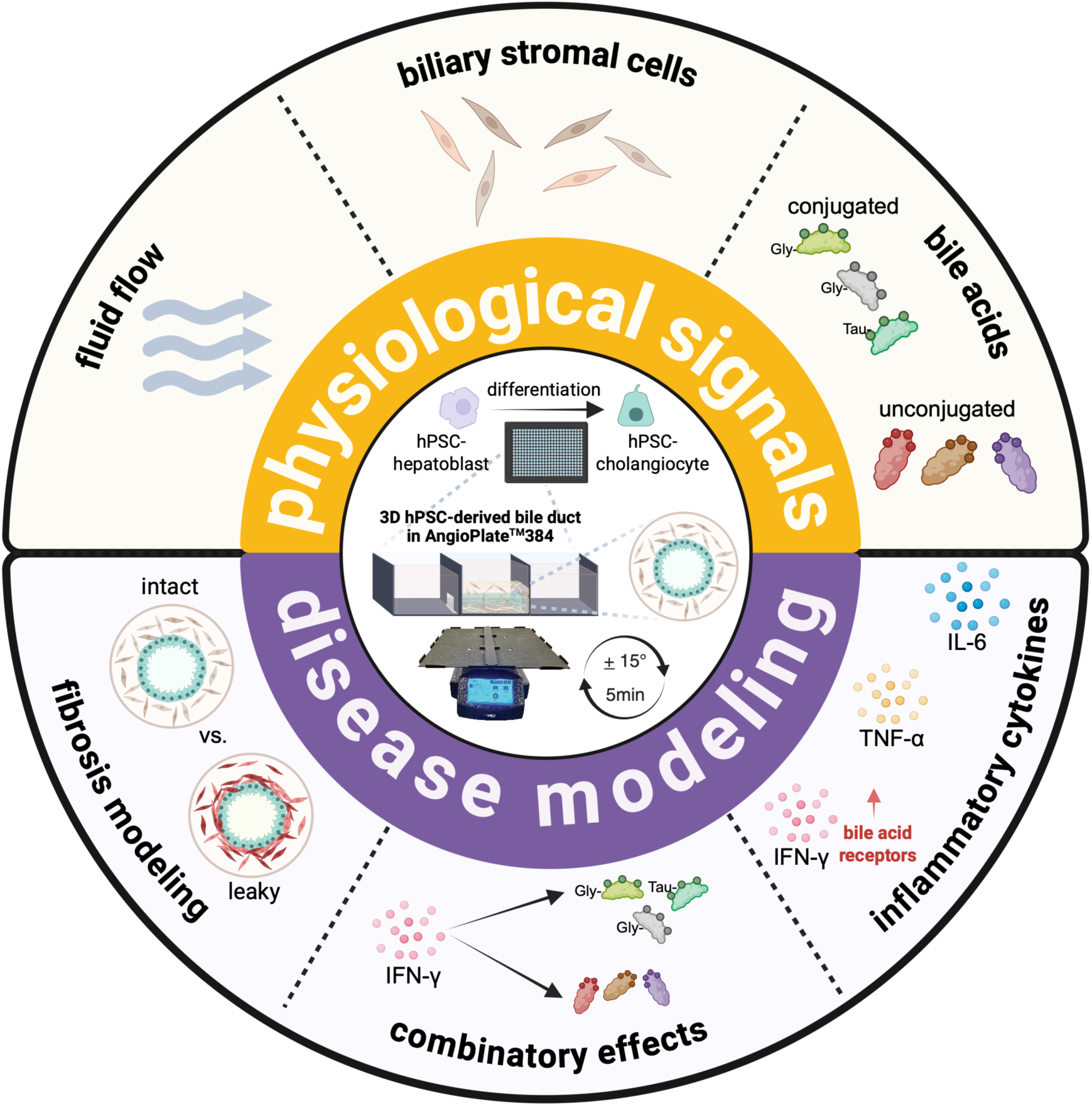

## Introduction

Despite recent advances in cholangiocyte organoid systems, no human cell-based platform yet faithfully recapitulates intrahepatic cholangiocyte physiology in vitro, limiting our ability to model complex biliary disease pathogenesis. Human induced pluripotent stem cell (hiPSC)-derived cholangiocyte organoids have been used for disease modelling, such as cystic dysfunction in Primary Sclerosing Cholangitis patient iPSC-derived extrahepatic organoids^[1]^, and biliary functional defects resulting from cystic fibrosis transmembrane conductance regulator (CFTR) mutations in cystic fibrosis patient iPSC-derived intrahepatic organoids^[2–4]^. Although these models can capture key aspects of biliary physiology and disease, they lack the open lumen of intrahepatic networks as see in vivo, restricting investigation into the more complex interplay between biliary constituents and fluid flow.

Intrahepatic cholangiocytes are continuously exposed to a concentrated stream of bile acids (BAs), which in humans are predominantly present as glycine-conjugated species with only trace levels of unconjugated forms^[5–7]^. BAs serve as critical niche-specific signals that shape cholangiocyte spatial identity, as demonstrated in a study by Sampaziotis et al. (2020) showing that human primary cholangiocyte organoids lost their native transcriptional signature when cultured in defined media but adopted a gallbladder-like identity when treated with gallbladder bile^[8]^. In disease contexts, the composition of the BA pool becomes dysregulated, as reflected by alterations detected in serum, fecal, and gallbladder analyses^[9–11]^. Yet, the effective intrahepatic BA concentrations and their specific physiological or pathological contributions remain poorly defined, highlighting the need for physiologically relevant in vitro platforms that can systematically interrogate how bile acid and flow dynamics influence intrahepatic cholangiocyte behavior in healthy and diseased states.

To overcome the limitations of conventional 3D organoid systems and enable simultaneous investigation of the pathological contributions of biliary constituents, advances in bile-duct-on-chip platforms offer a perfusable lumen that supports fluid flow and the controlled introduction of physiological biliary constituents. Fluid shear stress has been established as a key physiological regulator in these systems^[12,13]^. This field has also recently begun to explore cell-cell interactions, including endothelial cocultures with primary human cholangiocyte organoids derived from the extrahepatic bile duct of PSC patients to assess IL-17A-mediated inflammatory responses^[14]^. A recent study explored the formation of bile ducts with iPSC-derived hepatic progenitor cells co-cultured with immature mesenchymal and endothelial cells in a microfluidic platform; however, this system only recapitulates fetal liver signatures^[15]^, limiting its use for modelling more advanced chronic biliary diseases. While these studies underscore the importance of investigating cholangiocyte responses under physiologically relevant conditions, they mostly utilize murine cells or primary patient cells, and no study has yet systematically integrated these factors into a human intrahepatic cholangiocyte–based microfluidic platform to examine cellular responses. To enable the combinatorial testing of multiple factors that more accurately replicate the complex biliary environment, there is a growing need for improving the throughput of microfluidic devices over single-organ-on-chip systems.

To bright these gaps, we employed the AngioPlate™384 platform that can hold up to 128 perfusible hPSC-derived cholangiocyte tubes within a single plate^[16,17]^, with each middle well connected to inlet and outlet reservoirs to facilitate continuous fluid flow and an open-top design to allow easy tissue retrieval for downstream flow cytometry and molecular profiling. We showed that hPSC-cholangiocyte tube function is enhanced by fluid flow and stromal cell incorporation. Bile acid exposure, alone or in combination with a pro-inflammatory cytokine interferon-gamma, can modulate cholangiocyte responses. Lastly, we incorporated proliferative murine or hPSC-derived hepatic mesenchymal cells to model biliary fibrosis, a key feature of biliary disease.

## Results

### Efficient differentiation and epithelialization of 3D hPSC-chol tubes

Within the AngioPlate™384, tubular channels embedded in fibrin were formed through gravity-assisted seeding, enabling attachment, proliferation and expansion of hPSC-hepatoblasts (hPSC-HBs) along the luminal surface (**Fig. 1A**). Before seeding, hPSC-HBs were efficiently differentiated^[4,18]^ into a highly enriched population showing 98.3 ± 0.3% albumin (ALB) and alpha-fetoprotein (AFP) double-positivity (mean ± SEM) (**Supplementary Fig. 1A, B**). These progenitors displayed strong proliferative capacity and differentiation potential toward the cholangiocyte lineage. With modifications to our previous protocol^2^ (**Fig. 1B**), we achieved robust epithelialization of hPSC-chol tubes over 20 days (**Fig. 1C**), with 95.3 ± 0.6% of cells expressing cytokeratin 7 (CK7) on day 20 (mean ± SEM), confirming efficient cholangiocyte specification (**Supplementary Fig. 1C, D**). hPSC-chol maturation was further supported by the progressive downregulation of hepatic progenitor markers *ALB*, *AFP* and the maintenance of SRY-box transcription factor 9 (*SOX9*) cholangiocyte fate marker^[2,3,19,20]^, together with significant upregulation of the cholangiocyte maturation markers *CK7* and *CFTR* by day 20 (**Fig. 1D**).

**Figure 1.**
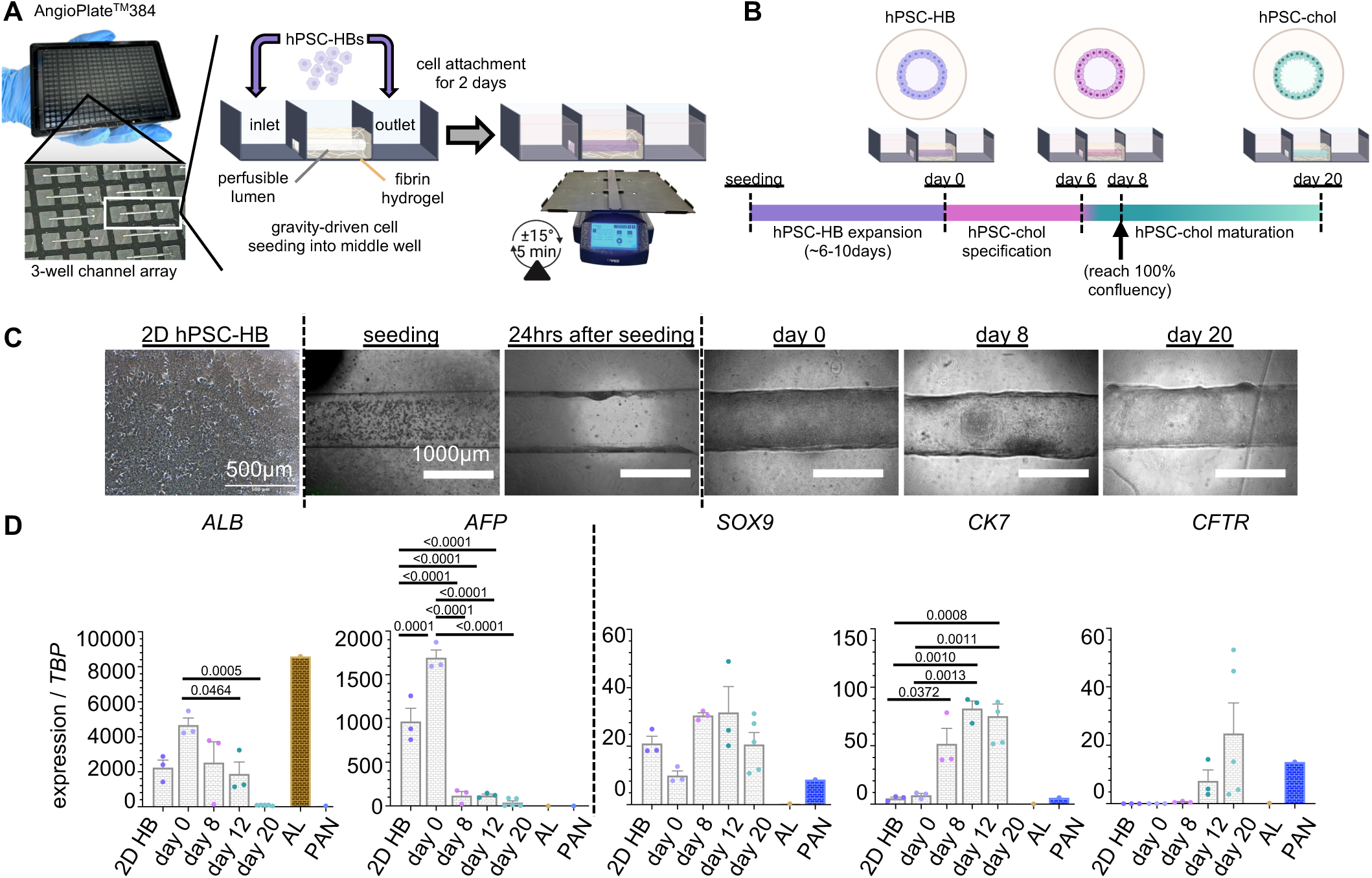
hPSC-chol tube differentiation in the 384-well AngioPlate. (**A**) Schematic for the assembly and working mechanism of the AngioPlate with gravity-based seeding process of hPSC-derived hepatoblasts (HBs). (**B**) hPSC-HB expansion and subsequent cholangiocyte tube differentiation with confluency maintained. (**C**) Representative images of the 3D hPSC-tube at each differentiation stage. (**D**) Quantitative PCR analysis of HB markers (*ALB, AFP*) and cholangiocyte markers (*SOX9, CK7, CFTR*) at each stage. *TBP*, TATA-box protein expression is set to 1-fold. AL, adult liver; PAN, adult human pancreas. Ordinary one-way ANOVA with Tukey’s comparison test performed on the first 5 groups including 2D HB and hPSC-chol tubes D0-20. N=3 experiments.

Apical-basal polarity is critical for cholangiocytes to mediate directional bile acid transport and maintain barrier integrity^[21,22]^. To validate cellular polarity, we performed immunostaining on day 20 hPSC-chol tubes. The apical primary cilium, crucial for fluid sensing and transport, was stained by α-tubulin and abundantly present as punctate signals along the luminal surface (**Fig. 2A**). Cilia appeared as apically oriented vertical projections extending into the lumen, positioned above the cell bodies visualized by CK7 and DAPI (**Fig. 2B**), with **Supplementary Video 1** further illustrating this point. The maturation marker CFTR co-localized with the tight junction marker ZO-1 (**Fig. 2C, D**), with both proteins exhibiting enriched expression at the apical surface, consistent with polarized epithelial organization (**Fig. 2D**). Collectively, these findings confirm the proper expression and localization of key functional maturation features in hPSC-chol tubes.

**Figure 2.**
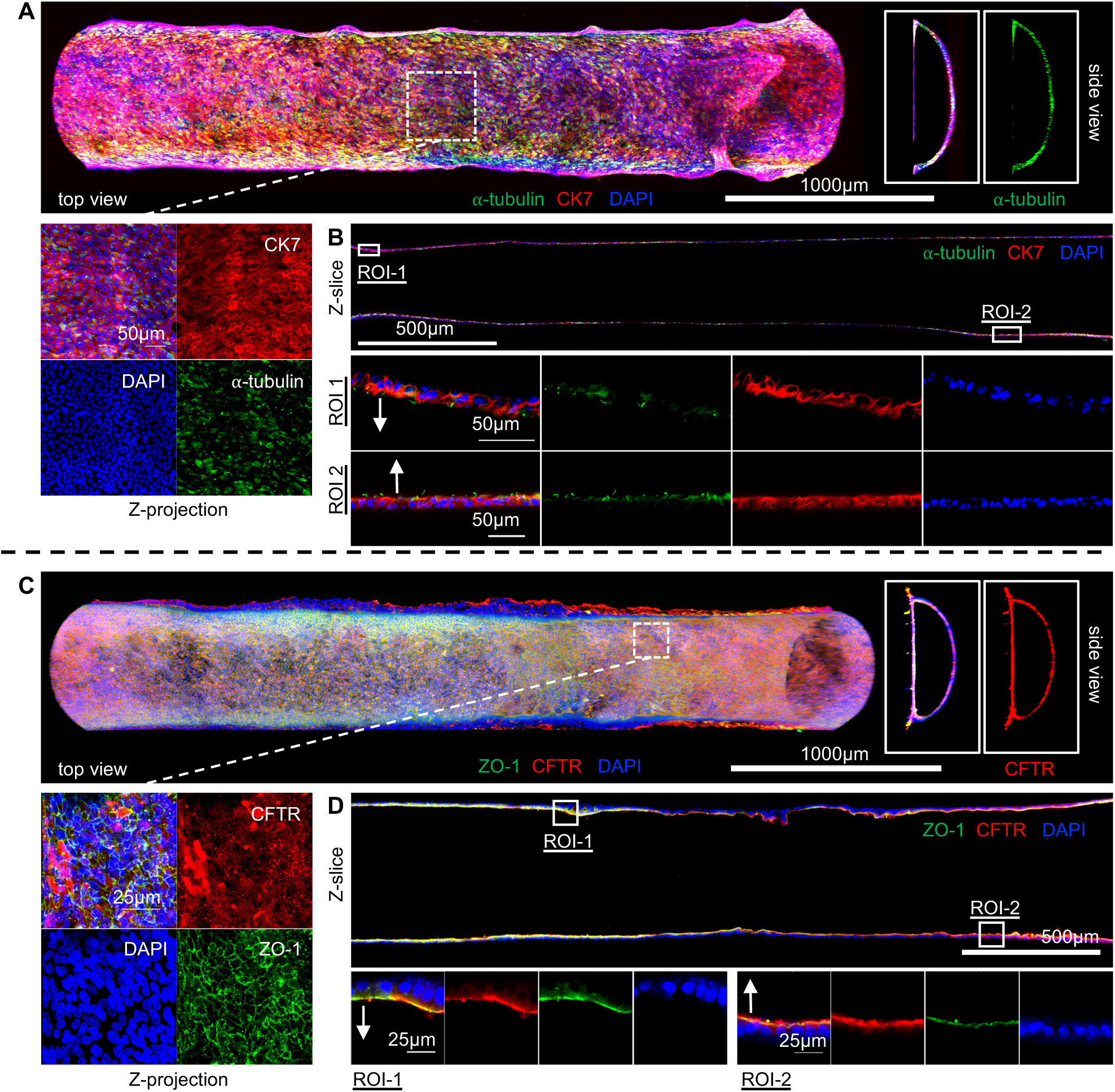
hPSC-chol tube exhibits apically polarized functional markers. (**A**) Immunostaining of CK7 and α-tubulin primary cilia marker. Enlarged Z-projection view showing CK7 and primary cilia distribution. Cilia are visible as bright α-tubulin specks. Image acquired in Resonant-mode (frame average 8); laser powers: DAPI 20.2; CK7 (red) 7.5; α-tubulin (green) 7.9. (**B**) Two regions of interests (ROIs) showing apically polarized cilia protruding towards the lumen. (**C**) Immunostaining of CFTR and ZO-1. Enlarged Z-stack projection view of CFTR and ZO-1 tight junction co-staining. Image acquired in Galvano-mode (exposure time 2.2 seconds; frame average 2); laser powers: DAPI 13.8; ZO-1 (green) 21.4; CFTR (red) 5.8. (**D**) Two regions of interest (ROIs) from a Z-slice of the tube showing apical enrichment of CFTR and ZO-1. For whole-tube renderings in panels (**A**) and (**C**), Non-linear LUT adjustments were made for better visualization of detailed features towards the top curvature of the tube. In panels (**B**) and (**D**), white arrows indicate the direction of the apical luminal space for each ROI.

### Fluid flow enhances hPSC-chol tube functional maturation

To further investigate the effect of fluid flow, we compared hPSC-chol tubes, which so far have been cultured on a rocker experiencing bi-directional fluid flow (flow), with those maintained under static conditions (static). As an initial readout, we assessed epithelial barrier function across three key stages of differentiation: day 0 (D0), when most HB tubes cultured under flow reached confluency and became ready for differentiation; day 8 (D8), corresponding to the end of retinoic acid-dependent cholangiocyte specification and initiation of ciliogenesis in maturation media; and finally, day 20 (D20), the maturation endpoint. Barrier integrity was assessed by perfusing tetramethylrhodamine-isothiocyanate (TRITC)-labelled dextran (70 kDa) and monitoring fluorescence leakage over time. A higher leakage rate indicates reduced epithelial integrity, visualized as red fluorescence diffusing into the surrounding hydrogel.

The barrier function improved over the 20-day differentiation period in both static and flow cultures, as indicated by progressively reduced luminal TRITC–dextran leakage (**Fig. 3A**), consistent with evidence that cholangiocytes acquire junctional markers as they mature^[21–23]^. Fluid flow further enhanced barrier integrity, with significantly lower leakage at all time points compared to static cultures (**Fig. 3A**). Flow also increased primary cilia formation in mature cholangiocytes from 27.4 ± 3.4% in static cultures to 45.9 ± 6.1% under flow (p=0.0289, mean ± SEM) (**Fig. 3B**). Correspondingly, the expression of two primary cilia markers, polycystin 1 (*PKD1*) and polycystin 2 (*PKD2*), also increased under fluid flow (**Fig. 3C**). *CFTR* gene expression was not affected by fluid flow (**Fig. 3D**). Since we previously detected primary cilia and CFTR apical polarization (**Fig. 2**), we assessed the expression of a planar-cell-polarity (PCP) regulator Van Gogh-Like 2 (*VANGL2*), that participates in biliary tree development^[24]^ and is especially important for primary cilia axoneme positioning and assembly across various cell types^[25,26]^. *VANGL2* was significantly upregulated under fluid flow (**Fig. 3E**). Meanwhile, another PCP gene, Van Gogh-Like 1 (*VANGL1*), which does not play as large a role as *VANGL2* in cholangiocyte PCP signalling^[24]^, was unchanged under fluid flow (**Fig. 3E**). Taken together, these results confirm that fluid flow enhances hPSC-chol functionality by strengthening epithelial barrier integrity and promoting ciliogenesis, both of which are tied to improved cellular polarity.

**Figure 3.**
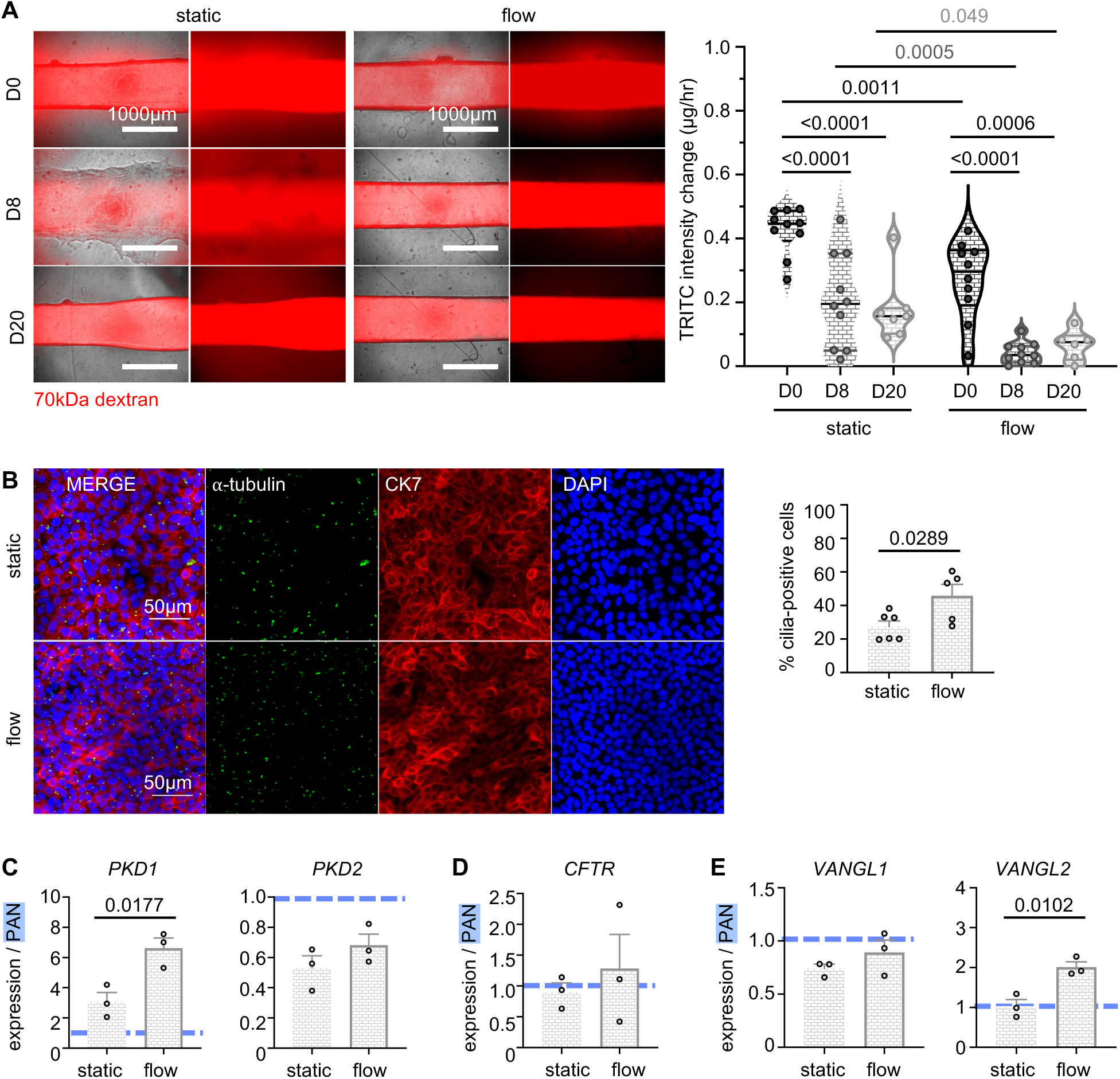
Fluid flow improves hPSC-chol tube function. (**A**) Representative fluorescent dextran leakage images and quantification of dextran leakage mass over time. N=3 experiments with 2-4 technical replicates per experiment. Two-way ANOVA with Tukey’s comparison test. (**B**) Confocal images of primary cilia (α-tubulin dots) representing one region of interest (ROI) in every hPSC-chol tube and corresponding cilia quantification. N=6 static experiments, N=5 flow experiments; unpaired T-test. Quantitative PCR analysis of (**C**) primary cilia markers *PKD1* and *PKD2,* (**D**) *CFTR,* and (**E**) planar cell polarity markers *VANGL1* and *VANGL2* mRNA expression levels. PAN, adult human pancreas gene expression is set to 1-fold and labelled with a dotted line for reference. N=3 experiments; unpaired T-test.

### Stromal cell incorporation enhances hPSC-chol functional maturation under fluid flow

The interaction between hepatic progenitor cells and periportal mesenchymal cells is critical for intrahepatic cholangiocyte specification and morphogenesis^[19]^. Beyond their role in fate specification, mesenchymal cells are also known to promote epithelial tight junctional formation^[27]^. To recapitulate this microenvironmental niche, we supported hPSC-derived cholangiocyte differentiation with OP9 stromal cells^[4]^ engineered to overexpress the Notch ligand Jagged-1 (OP9j). To modelling the biliary mesenchyme, irradiated OP9j cells (irrOP9j) were incorporated into the fibrin hydrogel (**Supplementary Fig. 2A**). hPSC-chol tubular differentiation in the presence of irrOP9j (**Supplementary Fig. 2B**) was then evaluated using the same quantity control parameters established for stromal-free tubes. IrrOP9j(+) tubes remained confluent throughout differentiation (**Supplementary Fig. 2C**) with *ALB* and *AFP* downregulation, *SOX9* persistence coupled with the simultaneous increase of *CK7* and *CFTR* maturation markers (**Supplementary Fig. 2D**). IrrOP9j(+) tubes also exhibited superior barrier function under fluid flow compared to static (**Supplementary Fig. 3A**). Fluid flow increased cilia-positivity to 70.3 ± 3.3% compared to 35.7 ± 2.6% in static cultures (p<0.0001, mean ± SEM) (**Supplementary Fig. 3B**), an improvement that was also further corroborated by enhanced expression of ciliary and PCP markers (**Supplementary Fig. 3C-E**).

As cilia positivity increased from ∼46% in stroma-free tubes to ∼70% with stromal cells under fluid flow, we directly compared the two conditions to confirm whether irrOP9j cells conferred any benefits. The addition of irrOP9j cells significantly increased barrier tightness (**Fig. 4A**) and, in agreement with previous results, improved primary cilia positivity from 42.5 ± 7.9% in flow-only tubes to 74.7 ± 4.7% (p<0.0001, mean ± SEM) (**Fig. 4B**). Correspondingly, flow irrOP9j(+) tubes showed higher expression of the ciliary markers *PKD1*, *PKD2* and polarity marker *VANGL2* (**Fig. 4C**). Confocal imaging of flow irrOP9j(+) tubes further confirmed apically polarized primary cilia (**Supplementary Fig. 4A**) and CFTR (**Supplementary Fig. 4B**). These results collectively suggest that, on top of fluid flow, stromal cells can further enhance the mature functional characteristics of hPSC-chol tubes by improving barrier function, ciliogenesis and polarity.

**Figure 4.**
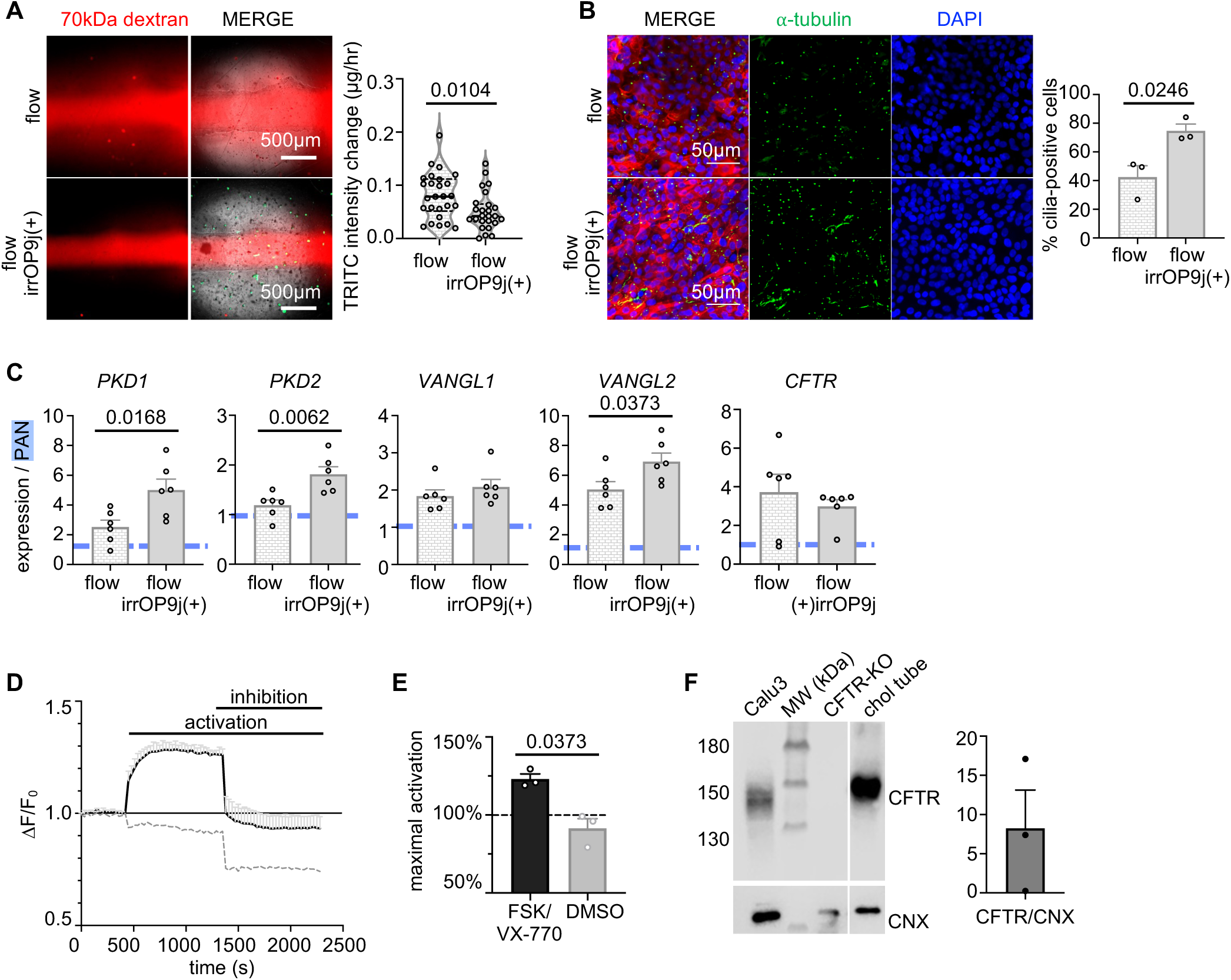
Stromal cells incorporation under fluid flow further enhances function in hPSC-chol tubes. (**A**) Representative images of TRITC 70kDa dextran leakage and quantification as mass over time. N=8 experiments with variable numbers of technical replicates; unpaired T-test. (**B**) Primary-cilia positivity quantification. N=3 experiments, unpaired T-test. (**C**) Quantitative PCR analysis of primary cilia markers *PKD1* and *PKD2,* the planar cell polarity markers *VANGL1* and *VANGL2,* and *CFTR* mRNA expression levels. PAN, adult human pancreas gene expression is set to 1-fold and labeled with dotted line for reference. N=3 experiments with two technical replicates per experiment; unpaired T-test. (**D**) Representative time-lapse plot from one FLIPR experiment (left), all fluorescence readings are normalized to the last timepoint of baseline reading that is set at 1.0 fold. Black line is FSK/VX-770 and grey line is DMSO. (**E**) The maximal point from the activation phase by either FSK/VX-770 or DMSO. N=3 experiments; unpaired T-test. (**F**) Representative Western Blot of CFTR protein expression from hPSC-chol tubes after the FLIPR assay and quantification of protein amount against Calnexin (CNX).

To further validate irrOP9j(+) hPSC-chol tubular function under fluid flow, we evaluated chloride conductance using the Fluorescence Imaging Plate Reader (FLIPR) assay^[28,29]^. Apical chloride transport, a core secretory function of cholangiocytes dependent on apical localization of the CFTR, was stimulated with the cyclic AMP agonist forskolin (FSK) and the CFTR potentiator ivacaftor (VX-770), resulting in an average maximal stimulation of 31.63 ± 0.06289% over DMSO activity (p=0.0373, mean ± SEM) (**Fig. 4D, E**). CFTR function was further corroborated by glycosylated CFTR protein expression detected via western blotting (**Fig. 4F)**. Taken together with the demonstrated epithelial polarity and barrier integrity, these findings indicate that our 3D hPSC-chol tubes recapitulated the key functional properties of human intrahepatic bile ducts.

### Increasing BA levels disrupted hPSC-chol epithelial integrity under fluid flow

To further model the biliary microenvironmental complexity, we treated hPSC-chol tubes with the six major human BAs: GCA, GCDCA, TDCA (conjugated BAs) and CA, CDCA, DCA (unconjugated forms), by estimating the low end of treatment ranges based on total BA pool mass and daily enterohepatic circulation^[19,30,31]^, while also accounting for receptor binding affinities^[31,32]^. BAs were added for 12 days during the cholangiocyte maturation to simulate chronic exposure.

Compared to control (ctrl), epithelial leakage was reduced significantly under treatment with low levels of GCDCA and TDCA (**Fig. 5A**), suggesting that they may exert a protective effect on cholangiocyte epithelium. At high levels, GCA 5.0mM induced significant leakage over control, while GCDCA 1.0mM and TDCA 0.5mM also induced visible barrier leakiness but did not reach statistical significance (**Fig. 5A**). Compared to conjugated BAs, the hydrophobic unconjugated BAs required much lower effective concentrations to elicit barrier disruption. Unconjugated BAs at low concentrations, similar to their conjugated counterparts, also show a trend of reducing barrier leakage; meanwhile, their corresponding alcohol vehicles did not exert a significant effect (**Supplementary Fig. 5A**). Among the high concentrations of unconjugated BAs, CDCA 0.25mM induced the highest barrier leakage (p<0.0001). (**Supplementary Fig. 5A**). These results suggest that low BA concentrations may support cholangiocyte barrier integrity while high concentrations damage the epithelium.

**Figure 5.**
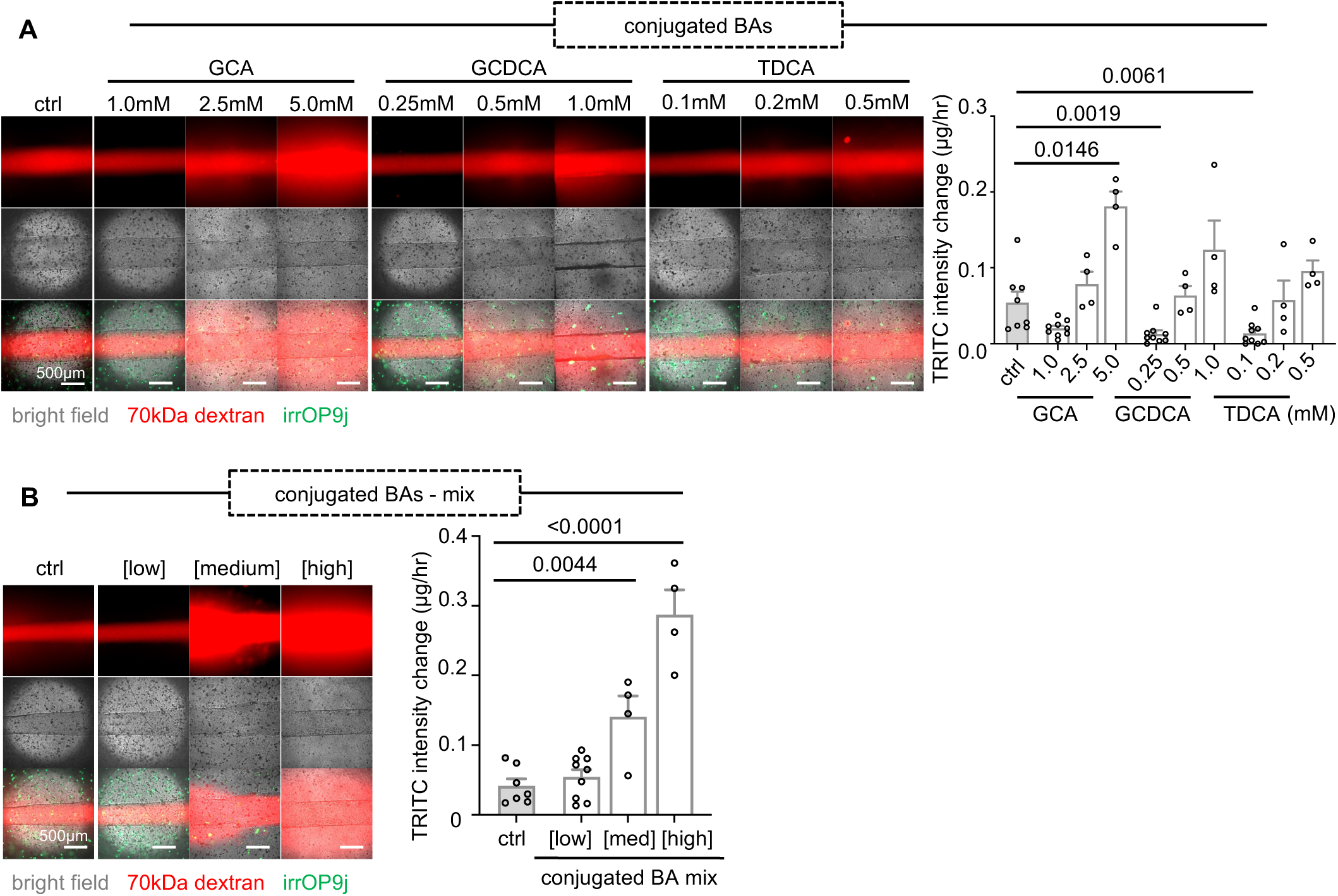
Effect of bile acid flow on hPSC-chol epithelial integrity. (**A**) 70kDa dextran leakage images of tubes treated with individual conjugated BA species from low, medium or high concentrations and leakage quantification mass over time. Lognormal ordinary One-way ANOVA. (**B**) Dextran leakage images of tubes treated with conjugated BAs mixed at low, medium or high concentrations and leakage quantification. Ordinary one-way ANOVA. N=3-4 experiments with 1-2 technical replicates per experiment. All ANOVAs are followed up with Dunnett’s test.

Rather than a single bile acid, human intrahepatic bile comprises a diverse BA mixture where their dynamic interactions critically influence cellular responses^[33]^. To better approximate this physiological complexity, we generated mixtures from the three dominant conjugated and unconjugated BAs at low, medium, and high concentrations. Increasing levels of both conjugated and unconjugated BA mixtures induced significant epithelial leakage (**Fig. 5B; Supplementary Fig. 5B**). Notably, cell death occurred after treatment with 0.25 mM CDCA and high BA concentration mixtures, whereas tubes remained intact without visible cell loss in all other conditions (**Fig. 5; Supplementary Fig. 5**). Together, these results revealed distinct thresholds at which BAs compromise epithelial integrity versus viability.

### Effects of inflammatory cytokines on hPSC-chol tubes

Besides bile acid accumulation, chronic biliary inflammation is a major driver of diseases impacting intrahepatic cholangiocytes, such as Primary Biliary Cholangitis^[34]^, Primary Sclerosing Cholangitis^[1,34,35]^ and Cystic Fibrosis Related Liver Disease^[36]^. Nevertheless, the direct effects of inflammatory cytokines specifically on human intrahepatic cholangiocytes in these diseases remain unclear. To explore this, we treated hPSC-chol tubes with three main pro-inflammatory mediators implicated in biliary diseases: interleukin-6 (IL-6), tumour necrosis factor-alpha (TNF-α), and interferon-gamma (IFN-γ). None of these cytokines caused barrier leakage (**Fig. 6A**). To see if other functional changes occurred without obvious barrier dysfunction, we analyzed the expression of various secretory transporter genes using quantitative PCR. IL-6 did not change the expression of the markers tested. 40 ng/ml TNF-α slightly increased the expression of the functional markers *CFTR*, *SCTR*, and *PKD2* (**Fig. 6B**). Compared to the other two cytokines, IFN-γ reduced all functional markers overall, with significant reductions in the two cholangiocyte chloride channels *CFTR* and anoctamin 1 (*TMEM16A*), as well as the ciliary marker *TRPV4* (**Fig. 6B**). This is further supported by primary cilia positivity declining from 76.7 ± 1.6% in ctrl to 50.2 ± 4.7% after treatment with 25 ng/ml IFN-γ (p=0.0060, mean ± SEM) (**Fig. 6C**). Notably, IFN-γ was the only cytokine to significantly increase the expression of two intrahepatic cholangiocyte bile acid receptors^[31]^: Takeda G-protein coupled receptor 5 (*TGR5*) and Farnesoid X receptor (*FXR*) (**Fig. 6D**).

**Figure 6.**
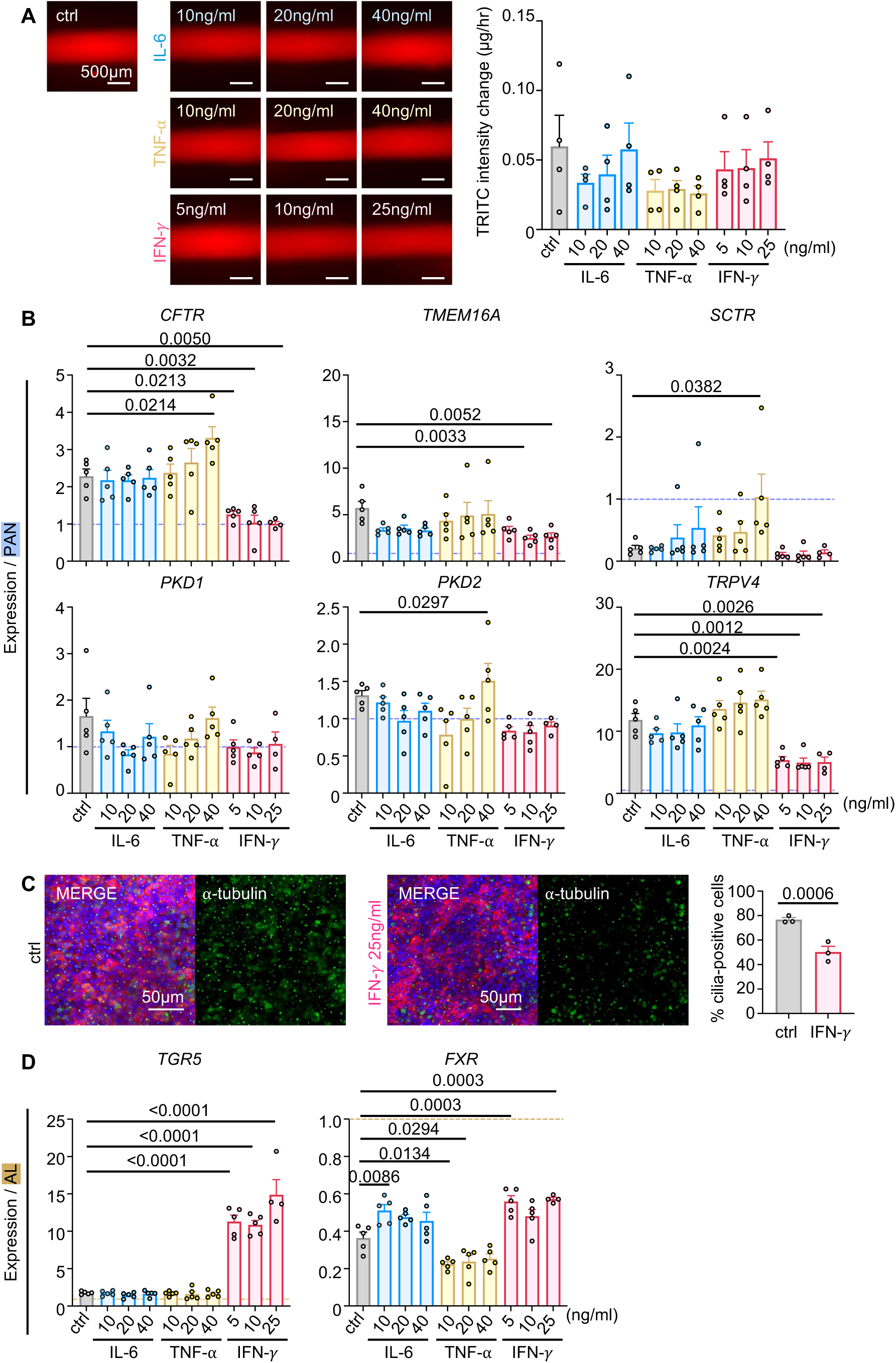
Effect of inflammatory cytokines on hPSC-chol tubes under fluid flow. (**A**) Representative fluorescent images of 70kDa TRITC dextran leakage status and dextran leakage quantification. Ordinary one-way ANOVA. (**B**) Quantitative PCR analysis of cholangiocyte functional markers after inflammatory cytokine treatment. PAN, adult human pancreas, is set to 1-fold and labeled with a dotted line. Ordinary one-way ANOVA. (**C**) Representative ROI stained with CK7 and primary cilia (α-tubulin) in ctrl or 25ng/ml IFN-γ treated hPSC-tubes. Primary cilia positivity quantification from N=3 experiments; unpaired T-test. (**D**) Quantitative PCR of bile acid receptor mRNA expression levels after inflammatory cytokine treatment. AL, adult human liver, is set to 1-fold and labeled with a dotted line. Panels (**A**), (**B**) and (**D**) are N=3 with 1-2 technical replicates per experiment. Ordinary one-way ANOVA or lognormal ordinary one-way ANOVA for each gene in panels (**B**) and (**D**). All ANOVAs are followed up with Dunnett’s comparison test.

### Combined effects of bile acids and inflammatory cytokines

Bile acids have been shown to activate immune cell signalling pathways downstream of TGR5 and FXR receptors^[31]^, but their roles in biliary inflammation are not clear. To this end, we assessed cholangiocyte response to IFN-γ in the hPSC-chol tubes without compromised viability after BA exposure. For conjugated BAs, IFN-γ exacerbated GCDCA-induced barrier leakiness in a dose-dependent manner, with the strongest effect observed at GCDCA 1.0 mM (**Fig. 7A; Supplementary Fig. 6A).** All medium and high concentrations of conjugated BAs reduced primary cilia, with more obvious reductions occurring at GCA 5.0mM, GCDCA 0.5mM, TDCA 0.2mM (**Fig. 7B; Supplementary Fig. 6B**). When combined with bile acid mixtures, IFN-γ had no additional impact on barrier integrity or primary ciliogenesis (**Supplementary Fig. 7**), and conjugated BA mixture at a medium concentration reduced primary cilia independent of IFN-γ (**Supplementary Fig. 7B**). Unlike with conjugated BAs, IFN-γ increased barrier leakage only in presence of low, but not high unconjugated BA concentrations (**Supplementary Fig. 8A**). Both CA 1.0 mM and DCA 0.1 mM independently reduced primary ciliation without additional effects from IFN-γ (**Supplementary Fig. 8B**). Together, these findings indicate that IFN-γ can synergize with specific BA species in a conjugation- and concentration-dependent manner to disrupt epithelial barrier function and ciliogenesis. This synergy emerged at higher concentration for conjugated bile acids, most prominently in combination with GCDCA, and at lower concentration for unconjugated BAs.

**Figure 7.**
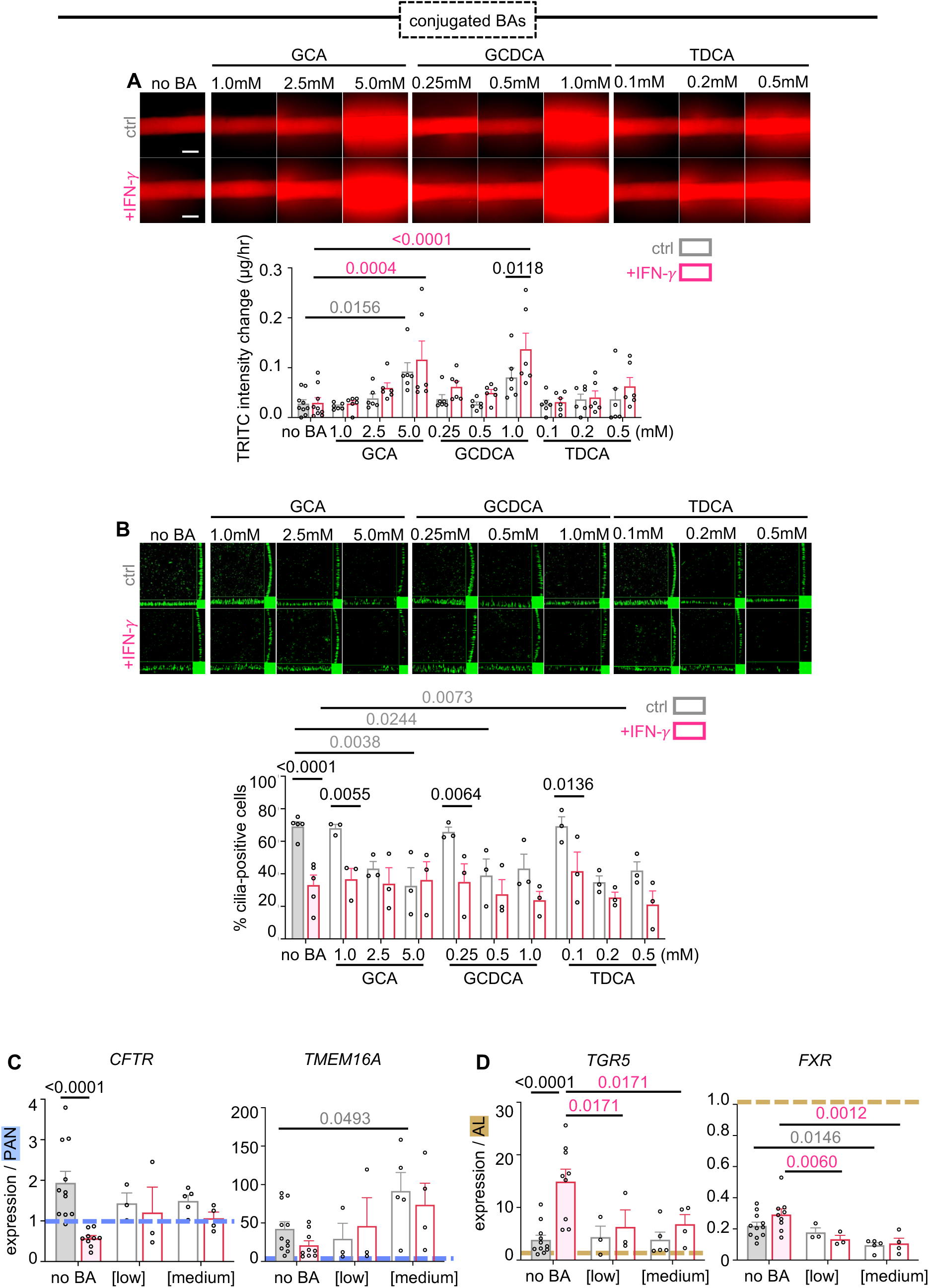
Conjugated bile acids can synergize with interferon-gamma to disrupt hPSC-cholangiocyte epithelium barrier integrity. **(A)** 70kDa dextran leakage representative images and quantification as mass-over-time for individual conjugated BA species and IFN-γ. N=3 experiments with 2 technical replicates for BA-treated groups and 3 technical replicates for no BA controls per experiment. (**B**) Primary cilia positivity representative images and quantification in hPSC-chol tubes treated by conjugated BAs and IFN-γ. N=3 experiments. Quantitative PCR analysis of (**C**) *CFTR* and *TMEM16A* chloride channel expression, and of (**D**) *TGR5* and *FXR* BA receptors in hPSC-chol tubes treated with conjugated BA mixtures and IFN-γ. N=3-5 experiments, two-way ANOVAs with Dunnett’s comparison test. Expression levels are normalized to PAN (adult human pancreas) for chloride channels or AL (adult human liver) for BA receptors, set to 1-fold and labelled with dotted lines.

Next, we examined the expression of key cholangiocyte chloride channels – CFTR and TMEM16A – by qPCR analysis to identify molecular alterations underlying the observed changes in response to BA and IFN-γ treatments. Individual conjugated BA species did not affect *CFTR* or *TMEM16A* expression (**Supplementary Fig. 9A**). As a mixture, however, medium levels of conjugated BAs increased *TMEM16A* expression (**Fig. 7C**). Low unconjugated BAs concentrations also minimally impacted functional gene expression, except for DCA 0.02 mM which reduced *CFTR* expression (**Supplementary Fig. 9B**), an effect that likely carried over into the low levels of unconjugated BA mixture (**Supplementary Fig. 8C**). High unconjugated BA concentrations downregulated *CFTR*, with IFN-γ further exacerbating this effect in presence of CA 1.0mM (**Supplementary Fig. 9B**).

Intriguingly, IFN-γ did not upregulate either *TGR5* or *FXR* in the presence of conjugated BA mixtures (**Fig. 7D**). Individual BA testing showed that low concentrations of GCA and GCDCA, as well as medium levels of TDCA suppressed IFN-γ-induced *TGR5* upregulation. Low GCA and TDCA also did so for *FXR* (**Supplementary Fig. 9C**). In contrast, unconjugated BAs, either as a mixture or individual species, did not significantly interfere with this upregulation (**Supplementary Fig. 8D; Supplementary Fig. 9D**). These results suggest that conjugated BAs can act in concert with IFN-γ to modulate bile acid receptor expression in cholangiocytes.

### Biliary fibrosis modelling using proliferative stromal cells in hPSC-chol tubes

Various hepatic mesenchymal cells, such as portal myofibroblasts, mesothelial and stellate cells, contribute to biliary injury signalling and fibrosis progression, during which they become activated, proliferate extensively and deposit extracellular matrix^[5]^. To model fibrosis within hPSC-chol tubes, non-irradiated OP9j cells were incorporated within the hydrogel. By day 20, these fibroblasts encased the lumen and induced substantial barrier leakage (**Fig. 8A**). Because murine cells have limited applicability for clinical-grade hepatobiliary disease modelling due to species-specific differences in hepatic metabolism and BA composition^[37]^, we replaced OP9j cells with hPSC-derived septum transversum mesenchyme (hPSC-STM), the progenitors of hepatic stromal lineages including portal fibroblasts and stellate cells^[38],[39]^. Like the OP9j, hPSC-STM cells also lead to a fibrotic phenotype and epithelial leakage (**Fig. 8A**). As a non-proliferative parallel to the irrOP9j cells, hPSC-STM were also irradiated (irrSTM(+) tubes), exhibiting similar barrier function as irrOP9j(+) hPSC-chol tubes (**Fig. 8A**).

**Figure 8.**
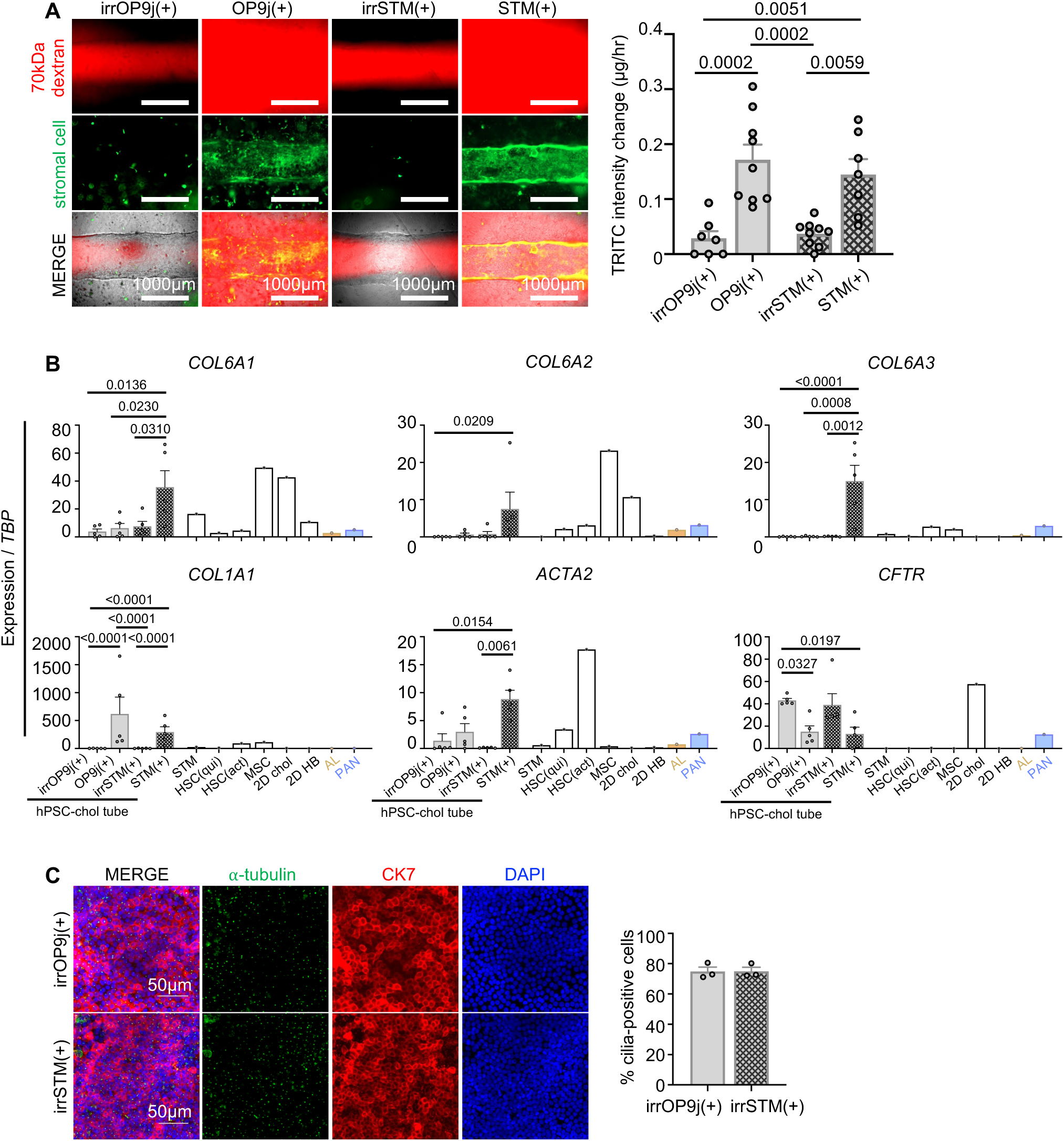
Hepatic stromal cells to support hPSC-chol tube function and model fibrosis. (**A**) Representative morphologies and 70kDa TRITC dextran leakage of day 20 hPSC-chol tubes cocultured with GFP-positive murine OP9j or hPSC-derived STM cells. (**B**) Quantitative PCR analysis of human pro-fibrotic genes and *CFTR* function marker in hPSC-chol tube cocultures. The following groups: STM, HSC (qui), HSC (act), MSC, 2D chol, 2D HB, AL and PAN are for reference and not included in statistical comparisons. Ordinary one-way ANOVAs with Tukey’s comparison test performed on the first 4 groups: irrOP9j(+), OP9j(+), irrSTM(+), and STM(+). N=3 experiments with 1-2 replicates per experiment. (**C**) Representative immunostaining of primary cilia (α-tubulin) and CK7 maturation marker, and cilia quantification.

Collagen type VI (*COL6A1*, *COL6A2*, and *COL6A3*), which show elevated biliary-specific deposition in liver fibrosis^[40]^, and two other general fibrosis markers, collagen type I (*COL1A1*) and alpha smooth muscle actin (*ACTA2*), were upregulated in hPSC-STM(+) tubes but not in tubes with irradiated STM or irradiated OP9j (**Fig. 8B**). *CFTR* expression was also reduced in cocultures with proliferative fibroblasts but remained high in irrOP9j(+) and irrSTM(+) cocultures (**Fig. 8B**). Primary ciliation was also comparable between irrOP9j(+) (74.9 ± 2.9%) and irrSTM(+) (75.0 ± 2.6%) hPSC-chol tubes under fluid flow (p=0.9704, mean ± SEM) (**Fig. 8C**). Overall, we showed that proliferative stromal cells drive fibrosis phenotypes by compromising barrier integrity, upregulating pro-fibrotic markers and downregulate CFTR. Additionally, hPSC-STMs can replace murine OP9j cells, serving either as a fibrosis model or, when irradiated, as supportive stroma that preserves cholangiocyte barrier function, gene expression, and primary ciliation at levels comparable to irradiated murine cells.

## Discussion

Our study comprehensively integrates a broad spectrum of physiologically relevant biliary signals ranging from fluid flow, stromal support, bile acids and inflammatory cytokines – leveraging a platform with markedly higher throughput than traditional bile-duct-on-chips. By layering the biliary signals of interest stepwise to reconstruct increasing biliary complexity, and subsequently superimposing pathological factors on this foundation, we uncovered divergent phenotypes and molecular signatures that may underlie distinct modes of intrahepatic cholangiocyte homeostasis or dysfunction.

We found that fluid flow enhanced cholangiocyte ciliogenesis and barrier integrity – two key features that sustain ion transport and bile modulation functionalities. These findings support the beneficial role of physiologically relevant fluid flow, consistent with prior evidence from other in vitro studies demonstrating improved barrier integrity in extrahepatic cholangiocytes under fluid flow^[12]^ and strengthened cellular junctions downstream of shear-stress-activated mechanosensing in vascular and neural systems^[14,41,42]^. For intrahepatic cholangiocytes, this flow response is mediated by the apical primary cilium, a single apical protrusion from the cell-body that bends in response to fluid stimuli^[4,43,44]^. The clinical relevance of primary cilia in cholangiocyte biology is further emphasized by recent evidence from Esser et al. (2024), who demonstrated that loss of primary cilia following ischemia-reperfusion injury in liver transplantation diminishes the biliary epithelium function and regenerative capacity that contributes to graft dysfunction^[45]^. In this context, our hPSC-derived 3D bile duct tubes can provide a powerful experimental platform: they not only maintain primary cilia under physiological flow but can also be adapted to model pathological conditions in which cilia are reduced or absent. This dual capability enables the interrogation of both normal biliary function and disease-relevant states, advancing our ability to study the mechanistic link between cilia integrity, cholangiocyte resilience, and regenerative potential in clinically relevant settings.

In examining bile acids (BAs), we identified distinct traits and molecular signatures in response to the dominant human conjugated and unconjugated BAs, particularly in their modes of interaction with the inflammatory cytokine IFN-γ. Conjugated BAs synergized with the cytokine to disrupt barrier integrity at higher concentrations, whereas for unconjugated BAs, this synergy appeared only at lower BA concentrations. Moreover, low concentrations of conjugated BAs suppressed IFN-γ-induced upregulation of the BA receptors *TGR5* and *FXR*, whereas unconjugated BAs did not. TGR5 is a known interferon-stimulated response element^[46]^, and our study is the first to demonstrate consistent type-II interferon–mediated upregulation of the pharmacologically relevant BA receptors, TGR5 and FXR, in hPSC-derived intrahepatic cholangiocytes. Prior studies showed that TGR5 activation by microbiota-derived DCA in dendritic cells^[47]^ as well as FXR activation in intestinal and hepatic macrophages^[48,49]^ suppress NF-κB signalling and mitigate tissue inflammation. While this mechanism has not been specifically confirmed for intrahepatic cholangiocytes; based on these reports, the BA receptor upregulation by IFN-γ could serve as a defence mechanism during the initiation phases of biliary disease. The suppression of this response by certain BAs may modulate these anti-inflammatory programs, alluding to distinct modes of cholangiocyte immune signaling depending on BA conjugation status. Future studies should delineate these complex cellular crosstalk dynamics in hPSC-chol tubular cocultures with immune cells under the context of BA-pool disruptions.

By dissecting the distinct pathological effects of bile acids and pro-inflammatory cytokines, our study identified experimental conditions that induced barrier leakiness without cell loss, recapitulating early disease stages in which viable cholangiocytes exhibit partial dysfunction marked by disrupted junctions, altered gene expression, and reduced ciliation. In contrast, exposure to highly concentrated bile acids caused substantial cell death, mimicking later stages of biliary disease characterized by toxic bile-mediated epithelial destruction^[50]^. Together, these models clarify how biliary signals drive disease progression and provide valuable insight into stage-specific disease phenotypes as well as the specific signals required to elicit them, thereby informing the development of future targeted therapeutic interventions.

In concordance with biliary mesenchymal support of cholangiocyte maturation and function through the supplementation of various growth factor and ligands^[19]^, we demonstrate a supportive role of stromal cell incorporation under fluid flow. Additionally, irradiated hPSC-derived STM showed potential to replace the murine-derived irradiated OP9j used for cholangiocyte differentiation^[4]^. In contrast, proliferative mesenchymal cells encircled the tube and induced epithelial barrier leakage, mirroring periductal fibrosis in vivo with disrupted epithelial homeostasis^[5]^. This cascade recapitulates the pathological progression of biliary diseases, in which advancing fibrosis compromises ductal integrity and lead to toxic bile leakage. By modeling both the onset and progression of fibrosis in vitro, this platform provides a versatile tool for dissecting disease mechanisms and testing therapies aimed at preserving or restoring biliary function.

Overall, our functional, ciliated hPSC-derived 3D cholangiocyte tubes represent a powerful platform to dissect how physiologically relevant signals preserve biliary epithelial homeostasis and how their disruption drives pathological change. By integrating models of both healthy and disease-associated conditions, this system mechanistically evaluates cholangiocyte function to inform the development of individualized strategies and provides a foundation for advancing precision medicine approaches in biliary disease.

## Author contributions

B.T. and S.O. contributed to the experimental design, data interpretation and manuscript writing. B.T. performed experiments, acquired data and conducted analyses. M.O. and M.K. supported the differentiation of hPSC-derived hepatoblasts (HBs) and septum transversum mesenchymal cells (hPSC-STM) and assisted with quantitative PCR analysis of activated and quiescent hepatic stellate cells (HSC). G.L. and L.H. from CB’s laboratory assisted with FLIPR assays, Western Blotting and analyzed the related data. F.Z., A.H. and S.D. from B.Z.’s laboratory contributed to the design and provision of the AngioPlate devices and provided technical assistance. M.O., C.B., and B.Z. provided conceptual guidance for experimental design and revised the manuscript. S.O. supervised the experiments, edited the manuscript, and approved the final version.

## Acknowledgements

We gratefully acknowledge members of the Ogawa, Zhang and Bear laboratories for their valuable input on experimental design and manuscript preparation. We thank Dr. Feng Xu from the Advanced Optical Microscopy Facility (AOMF) for providing microscopy training and assistance. Illustrations in this publication were created using BioRender. This work was supported in part by the Medicine by Design Initiative at the University of Toronto, funded through the Canada First Research Excellence Fund (CFREF), the Canadian Institutes of Health Research (CIHR; Grant IDs: 1023968 and 1021677); and JSPS KAKENHI (Grant number 18K08589) awarded to S.O. Additional support was provided by the Stem Cell Network through an Impact Award (Grant ID: 1026927) awarded by S.O and B.Z. B.T. was a recipient of the 2023 Canadian Graduate Scholarships – Master’s (CGS-M) supported by the Canadian Institutes of Health Research.

## Competing interests

The AngioPlate^TM^384 is commercialized by OrganoBiotech, Inc. B.Z. is the co-founder and hold equity in the company. F.Z. and S.D. are also part-time employees of OrganoBiotech. The other authors declare that they have no competing interests.

## Materials and Methods

### AngioPlate gel casting

The AngioPlate was fabricated as previously described^[16,17]^, using gelatin as a sacrificial ink to pattern tubular structures spanning every three wells. AngioPlate (A002) is also commercially available from OrganoBiotech. Fibrin hydrogel was prepared by mixing fibrinogen (Sigma-Aldrich, F3879) with 7 U/ml thrombin (Sigma-Aldrich, T6884) in DPBS without calcium or magnesium (DPBS-/-, Corning, 21-031-CV) at a 5:1 ratio. A 25 μL fibrin droplet was added to each mid-well, and the plate was tapped rapidly 3–5 times for full coverage of the sacrificial structure. The plate was incubated statically at 37 °C for 30 min to allow fibrin crosslinking and gelatin degradation. Wells were then filled with DPBS-/- + 0.1% bovine serum albumin (BSA, Sigma-Aldrich, A1470) or stromal cell media (if stromal cells were used) and placed onto a rocker for 30-60 min at 37 °C to initiate luminal perfusion. Afterwards, tubes were washed twice with DPBS-/- + 0.1% BSA and rocked for an additional 20 min at 37 °C to remove excess gelatin. For luminal coating, 200μl of IMDM with 2.5% Matrigel (Corning, 354230) and 0.2 mg/ml aprotinin (Sigma-Aldrich, 616370) was added per tissue. When stromal cells were embedded in the hydrogel, the coating medium additionally contained 10 μM Y-27632 (Tocris, 1254) to promote survival. Plates were rocked overnight at 37 °C. Suitable rockers include the MIMETAS OrganoFlow® and OrganoBiotech IFlowRocker™.

### hPSC-derived hepatoblast progenitor preparation

H9 (WAe009-A) hPSC-HBs were generated as described by our group^[18]^. Passage-1 HBs from cryopreserved HB vials were used for AngioPlate cholangiocyte differentiation. Before seeding, 60μl expansion medium was added on top of the gel for hydration. Dissociated HBs were resuspended in expansion medium with 10μM Y-27632 and 0.5mg/ml aprotinin at 2 × 10⁶ cells/ml, and 100μl of cells was dispensed into each inlet/outlet. Plates were kept static for 2 days to allow attachment, then placed under continuous bi-directional fluid flow (15° tilt, 5-minute direction switching intervals), or kept flat throughout the expansion and differentiation period if it was a static culture. Media was changed every 2 days with freshly added 0.5 mg/ml aprotinin.

### hPSC-derived cholangiocyte tubular differentiation

The cholangiocyte backbone medium contained DMEM/F12 (Corning, 10-092-CV), 0.1% BSA, insulin-transferrin-selenium (Wisent, 315-082-QL), 1% penicillin-streptomycin (Wisent, 450-201-EL), 1% N21-MAX vitamin A-free media supplement (R&D Systems, AR012), 5% knockout serum (Gibco, 10828-028), 2 mM glutamine (Thermo Fisher, 25030081), 50 μg/ml L-ascorbic acid (Sigma-Aldrich, A4544), and 4.5 μM monothioglycerol (Sigma-Aldrich, M6145). Media were changed every two days with freshly added 0.5 mg/ml aprotinin. hPSC-HBs were specified toward cholangiocytes with 2.0μM retinoic acid (Sigma-Aldrich, R2625) for 6 days, during which cultures received 6 μM SB-431542 (Tocris, 1614) for the first 2 days, then 0.2 μM LDN-193189 (Tocris, 6053) and 5 μM forskolin (Tocris, 1099) for the next 4 days to prevent cell loss. Maturation was continued with LDN, FSK, and Y-27632 (Tocris, 1254) until day 20.

### Dextran permeability tests

Tubes were perfused with 200 μl of 0.1 mg/ml tetramethylrhodamine isothiocyanate-dextran (Sigma-Aldrich, T1162) in DPBS-/- + 0.1% BSA added to each inlet/outlet, with 60 μl of DPBS-/- + 0.1% BSA on top of the gel. For day 0 and day 8 measurements, tubes were perfused on the rocker for 1 hour. At day 20, perfusion was extended to 3 hours to increase sensitivity and reduce negative readings after background subtraction. Fluorescence was measured on either the Agilent CYT5M (Gen 5.3.4 software) or Fluostar Omega (MARS data analysis software) plate reader, and changes in intensity were calculated by readings at endpoint (1h or 3h) minus 0 min, normalized to a standard dilution curve^[16]^.

### Stromal cell preparation

OP9j cells were cultured in 80% minimum essential medium alpha medium (Gibco, 12000-022), 20% fetal bovine serum (FBS, Wisent, 090150) and 2mM glutamine as previously described^[51]^. GFP hPSC-derived STMs were differentiated from HES2-ESCs following a published protocol to generate WT-1 positive mesoderm cells^[38],[52]^ with modification (manuscript under preparation). OP9j and STM cells were irradiated in a Gamma Cell™ 3000 Elan Cesium-137 irradiator at 30 Gy. Following irradiation, the OP9j were frozen in 50% minimum essential medium, 40% FBS and 10% DMSO (Sigma-Aldrich, D2650) at -80 °C. IrrOP9j frozen vials were used in all experiments. Murine OP9j (non-irradiated and irradiated) were added at a density of 12K cells per fibrin gel, while for hPSC-STM (non-irradiated and irradiated), 1K cells were added per fibrin gel. Irradiated hPSC-STM cultures required 2mg/ml aprotinin for coating and culture media changes.

### Immunostaining

Tubes were perfused with 100µl 4% paraformaldehyde (Electron Microscopy Science, 15710-S) in DPBS with calcium and magnesium (DPBS+/+, CORNING, 21-030-CM) to fix samples for 15-20 minutes. The tubes were then washed with 0.1% BSA in DPBS+/+ (100µl per inlet/outlet and 60µl in mid-well) and rocked gently a few times. Washes were repeated 3 times at 5-minute intervals before and after all permeabilization and antibody-addition steps. CFTR, CK7 and ZO-1 proteins required 100% cold methanol permeabilization at 4 °C for 10 minutes. The staining steps that followed are as described in our previous publication^[4]^. Antibodies are listed in **Supplementary Tables 1 and 2**.

### Flow cytometry analysis

Cells were fixed in 4% paraformaldehyde in DPBS-/- for 15-20 minutes at room temperature. Cells were permeabilized with 90% cold methanol for 10 minutes at 4 °C. Human (ALB), alpha-fetoprotein (AFP) and cytokeratin 7 (CK7) flow cytometry was performed as previously described^[4]^. For day 20 hPSC-chol tubes,100μl of 1U/ml dispase (STEMCELL Technologies, 07923) was added to every tube to digest the fibrin gel for 35 minutes, followed by 5-10 minutes TrypLE™ Express (Gibco, 12605-010) treatment to break into single cells before fixation. All cells were filtered through a 70-100 μm nylon mesh to prevent clogging prior to running. Experiments were performed on a BD Biosciences LSRFortessa flow cytometer and analyzed using the FlowJo 10 software. 5000-10,000 total events were acquired in every experiment. Antibodies are listed in **Supplementary Tables 1 and 2.**

### Microscopic imaging

Live-cell fluorescent and brightfield images for dextran permeability tests were captured using either the Zeiss AxioObserver.Z1 or the EVOS FL2000 (Life Technologies) microscope. Whole-tube and primary cilia images were captured using Z-stacks with the Nikon A1R Confocal microscope at 20x magnification. Tiles were stitched using 10% blending for images spanning the entire tube (**Fig.2 and Supplementary Fig.4**). Whole-tube 3D volume rendering in **Fig.2A** was done in the NIS-Elements software using maximum-intensity display. For display purposes only, non-linear LUT adjustments were applied uniformly to the entire structure to increase visibility of the detailed features along the curved regions of the tube.

### Primary cilia quantification

All confocal images used for counting and display purposes in figures are Z-stack maximum intensity projections made in NIS-Elements. Each datapoint equals the number of α-tubulin-positive protrusions divided by nuclei (DAPI), counted within a 150μm×150μm central area in an ROI, averaged over 3 ROIs per experiment. Object counting was done using the ‘cell-counter’ feature in ImageJ software.

### FLIPR and Western Blotting assays

hPSC-chol tubes were cultured for 25 days to achieve stable CFTR expression. Tubes were digested with dispase (1U/ml) for 35 minutes, followed by TrypLE addition for 2-3 minutes to break the tubes into smaller pieces. Cholangiocytes from 5-8 tubes were combined into a single well of a 96-well black plate (CORNING, 3603). Cholangiocyte maturation medium was changed daily for 3 days with 10µM Y-27632 for the first day, and forskolin was removed 24 hours before the assay. Wells were loaded with a membrane potential dye (BLUE-R8034, Molecular Devices) dissolved in sodium-chloride-free buffer containing 150mM Gluconate lactone (A13105, Alfa Aesar), 150mM NMDG (66930, Sigma-Aldrich), 10mM HEPES (H3375, Sigma-Aldrich) and 3mM Potassium gluconate at pH 7.35 and 292 mOsm (G-4500, Sigma-Aldrich) for 30 minutes at 37 °C. The plate was then read in a fluorescence plate reader, SpectraMax i3 (Molecular Devices) at 37 °C. After the 7-minute baseline reading, CFTR was stimulated using 10µM forskolin (F6886, Sigma-Aldrich) ± 1µM VX-770 (S1144, Selleck Chemicals). After a 15-minute stimulation read, 10µM CFTR inh-172 (Cystic Fibrosis Foundation Therapeutics) was added to inhibit CFTR. The peak changes in fluorescence to CFTR agonists were normalized relative to fluorescence immediately before the addition of forskolin. After the assay, FLIPR dye was removed, and the plates were stored at -20°C for Western Blotting. Wells were lysed with lysis buffer (50 mM Tris-HCl, 150 mM NaCl, 1 mM EDTA, pH 7.4, 0.2 % (v/v) SDS, 0.1 % (v/v) Triton X-100, 1X inhibitor cocktail), after which Western Blotting was performed as previously described^[53]^.

### Quantitative PCR analysis

AngioPlate tubular structures were dissociated with 1U/ml dispase as described above. 100 μl lysis buffer was added to dissolve cell pellets for RNA purification using the Aqueous™-Micro Total RNA Isolation Kit (Invitrogen, AM1931). 200-1000 ng of RNA was mixed with iScript™ Reverse Transcription Supermix (BIO-RAD, 1708841) to make complementary DNA. Quantitative PCR was performed on a CFX96 Touch Real-Time PCR Detection System, with SsoAdvanced Universal SYBR® Green Supermix (BIO-RAD, 1725274). Bulk RNA samples for adult human liver (Clonetech, normal livers pooled from 3 male Asians of 22-64 years old, product number 636531, lot number 1402003) and adult human pancreas (Clonetech, normal pancreas from a 35-years old Caucasian, product number 636577, lot number 1703157A) were used as reference controls. Oligonucleotide sequences are listed in **Supplementary Table 3**.

### Bile acids and inflammatory cytokines

Bile acids were treated from day 8 to 20. Glycocholic acid (Sigma-Aldrich, G7132), glycochenodeoxycholic acid (Sigma-Aldrich, G0759), and taurodeoxycholic acid (Sigma-Aldrich, T0875). Cholic acid (Sigma-Aldrich, C1129) was reconstituted in methanol. Chenodeoxycholic acid (Cayman chemicals, 10011286) and deoxycholic acid (Sigma-Aldrich, D2510) were both reconstituted in ethanol (GREENFIELD, 22734-P006-EAAN). Inflammatory cytokines were treated from day 18 to 20. Interferon-gamma (IFN-g, PeproTech 300-02), tumor necrosis factor-alpha (TNF-a, proteintech, HZ-1014) and interleukin-6 (IL-6, R&D, 206-IL).

### Statistical Analysis

Each dataset includes at least three biological replicates (N), with each N representing an hPSC-chol tube differentiation from a single cryopreserved hPSC-HB vial. Data were analyzed using GraphPad Prism 10, with p < 0.05 considered significant and error bars representing mean ± SEM. Two-sample comparisons used two-tailed unpaired t-tests. For one-way ANOVAs, datasets were first tested for normality using the D’Agostino-Pearson omnibus (K2) test; non-normal datasets were assessed for lognormality and, if passing, analyzed with lognormal ordinary one-way ANOVA. Datasets failing both normal and lognormal assumptions were analyzed by the Kruskal-Wallis test with Dunn’s correction. Two-way ANOVAs included the Geisser-Greenhouse correction for sphericity. Tukey’s test was used for comparisons between all group means, and Dunnett’s test for comparing multiple treatments to a single control. For each figure, the N numbers and type of statistical tests employed were described in all figure legends.

## Notes

### Competing Interest Statement

The AngioPlateTM384 is commercialized by OrganoBiotech, Inc. B.Z. is the co-founder and hold equity in the company. F.Z. and S.D. are also part-time employees of OrganoBiotech.

