## Supplementary Text for "Bioengineered 3D hPSC-cholangiocyte ducts with physiological signals for biliary disease modelling"

**Supplementary Figure 1. Purity of starting hPSC-hepatoblast and ending hPSC-cholangiocyte populations in the AngioPlate.** (**A**) Example gating strategy for flow cytometry analysis of albumin (ALB) and alpha-fetoprotein (AFP) double-staining on the hPSC-HBs used for in-tube cholangiocyte differentiation. (**B**) Bar graphs of the population proportions in each quadrant from the contour plot from panel (A). N=27 hPSC-HB experiments; ordinary one-way ANOVA with Tukey's comparison test. (**C**) Example gating strategy for cytokeratin 7 (CK7) cholangiocyte marker in day 20 hPSC-chol tubes. (**D**) Bar graph of the CK7+ portion. N=4 hPSC-chol tube differentiations in the AngioPlate.

**Supplementary Figure 2. HPSC-chol tube differentiation with stromal cells in the fibrin hydrogel.** (**A**) Schematic of AngioPlate fibrin gel casting with irradiated murine OP9-jagged 1 cells (irrOP9j) incorporation. (**B**) The differentiation protocol while maintaining confluency in presence of irrOP9j. (**C**) Representative images of the irrOP9j(+) 3D hPSC-tube at each differentiation stage. (**D**) Quantitative PCR analysis of HB markers (*ALB, AFP*) and cholangiocyte markers (*SOX9, CK7, CFTR*) at each stage. *TBP*, TATA-box binding protein expression is set to 1-fold. N=3 experiments; ordinary one-way ANOVA with Tukey's comparison test.

**Supplementary Figure 3. The effect of fluid flow on hPSC-chol tubes in the presence of stromal cells.** (**A**) Representative fluorescent dextran leakage images and quantification of dextran leakage mass over time. N=3 experiments with 2-4 technical replicates per experiment; two-way ANOVA with Tukey's comparison test. (**B**) Confocal images of primary cilia (α-tubulin dots) representing one region of interest (ROI) in every hPSC-chol tube (fluorescent images) and corresponding primary cilia quantification (bar graph). N=6 static experiments, N=5 flow experiments; unpaired T-test. Quantitative PCR analysis of the mRNA expression levels of (**C**) primary cilia markers *PKD1* and *PKD2,* (**D**) *CFTR,* (**E**) planar cell polarity markers *VANGL1* and *VANGL2*. PAN, adult human pancreas gene expression is set to 1-fold and labelled with a dotted line for reference. N=3 experiments; unpaired T-test.

**Supplementary Figure 4. Apical basal polarity in hPSC-chol tubes with stromal cells.** (**A**) Z slice of CK7 and α-tubulin primary cilia marker immunostaining, with two regions of interests (ROIs) from a Z-slice of the tube, showing apically polarized cilia protruding towards the lumen. (**B**) Z slice of CFTR and ZO-1 immunostaining with two regions of interest (ROIs) showing apical enrichment of the two proteins. In ROIs, white arrows indicate the direction of the apical luminal space.

**Supplementary Figure 5. Unconjugated BAs and alcohol vehicle tests**. Representative brightfield and 70kDa dextran leakage images and quantification, in tubes treated with unconjugated BAs and their respective alcohol vehicles as (**A**) individual components, N=4 experiments, or (**B**) mixed at low, medium or high concentrations, N=5 experiments. CA was reconstituted in methanol (MeOH), CDCA and DCA in ethanol (EtOH). Ordinary one-way ANOVA with Dunnett's comparison test.

**Supplementary Figure 6. Barrier integrity tests and primary cilia positivity in bile acid and interferon-gamma combinatory treatments.** Representative images of (**A**) 70kDa dextran leakage and (**B**) primary cilia, CK7 and DAPI immunostaining in hPSC-chol tubes treated with individual conjugated BA species and IFN-$\gamma$.

**Supplementary Figure 7 Treatment of hPSC-chol tubes with bile acid mixtures and interferon-gamma.** (**A**) Representative 70kDa dextran leakage images and quantification, N=3 experiments; and (**B**) representative primary cilia, CK7 and DAPI immunostaining and cilia quantification in hPSC-chol tubes treated with conjugated BA mixtures and IFN-$\gamma$. N=3 experiments. Two-way ANOVA with Dunnett's comparison test for (**A**) and (**B**). (**C**) Representative 70kDa dextran leakage images and quantification, N=9 experiments, and (**D**) representative primary cilia, CK7 and DAPI immunostaining and cilia quantification, in hPSC-chol tubes treated with unconjugated BA mixtures and IFN-$\gamma$, N=3 experiments. Two-way ANOVA with uncorrected Fisher's LSD test for (**C**) and (**D**). For experiments testing BA mixtures alongside individual species, the 'no BA' control datapoints for conjugated mixtures correspond to those shown in Fig. 7A (dextran leakage assay) and Fig. 7B (primary cilia), while the 'no BA' control datapoints for unconjugated mixtures correspond to those in Fig. 7C (dextran assay) and Fig. 7D (primary cilia).

**Supplementary Figure 8. Combinatory effects of unconjugated bile acids and interferon-gamma on hPSC-chol tube response.** (**A**) 70kDa dextran leakage representative images and quantification as mass-over-time for individual unconjugated BA species from low to high concentrations and IFN-$\gamma$. N=6-8 experiments. (**B**) Primary cilia positivity representative images and quantification in hPSC-chol tubes treated by unconjugated BAs and IFN-$\gamma$. N=3 experiments.

Quantitative PCR analysis of (**C**) *CFTR* and *TMEM16A* chloride channel expression, and of (**D**) *TGR5* and *FXR* BA receptors in hPSC-chol tubes treated with unconjugated BA mixtures and IFN-$\gamma$. N=6 experiments, two-way ANOVAs with uncorrected Fisher's LSD test. Expression levels are normalized to PAN (adult human pancreas) for chloride channels or AL (adult human liver) for BA receptors, set to 1-fold and labelled with dotted lines.

**Supplementary Figure 9 Quantitative PCR analysis of functional gene expression in hPSC-chol tubes treated with single bile acids.** Chloride channel *CFTR* and *TMEM16A* expression levels in (**A**) conjugated BA-treated tubes and (**B**) unconjugated BA-treated tubes. BA receptor *TGR5* and *FXR* expression levels in (**C**) conjugated BA-treated tubes and (**D**) unconjugated BA-treated tubes. N=4-6 experiments for conjugated BAs, and N=3-5 experiments for unconjugated BAs. Two-way ANOVA with Dunnett's comparison test for all graphs.

**Supplementary Figure 10 Uncropped Western Blot gel image.** Boxes indicate the bands displayed in main **Fig. 4E**. Arrows indicate the bands used for quantification in main **Fig. 4F**.

**Supplementary Video 1 A walk through the 3D hPSC-chol tube luminal space**. The tube is the same as the one displayed in main Fig. 2. Primary cilia marked by α-tubulin (in green) are seen protruding from the surface of the tube inwards towards the luminal side with exposure to fluid flow.

### Supplementary Table 1. Primary antibody list.

| Name | Vendor | Product number | Host and specificity | Dilution |
|---|---|---|---|---|
| CFTR (24-1) | R&D Systems | MAB25031 | Mouse, anti-human | 1:200 |
| CFTR (13-1) | R&D Systems | MAB1660 | Mouse, anti-human | 1:200 |
| CK7 | Abcam | AB68459 | Rabbit, anti-human | 1:200 |
| Acetylated alpha-tubulin | Sigma Aldrich | T7451 | Mouse, anti-human | 1:800 |
| ZO-1 | Thermo Fisher | 40-2200 | Rabbit, anti-human | 1:400 |
| ALB | Bethyl | A80-129A | Goat, anti-human | 1:200 |
| AFP | DAKO | A0008 | Rabbit, anti-human | 1:2000 |

### Supplementary Table 2. Secondary antibody list.

| Name and target species | Vendor | Product number | Dilution |
|---|---|---|---|
| Donkey anti-Mouse Alexa488 | Invitrogen | A21202 | 1:400 |
| Donkey anti-Rabbit Alexa555 | Invitrogen | A31572 | 1:400 |
| Donkey anti-Mouse Alexa555 | Invitrogen | A31570 | 1:400 |
| Donkey anti-Rabbit Alexa488 | Invitrogen | A21206 | 1:400 |
| Donkey anti-Mouse Alexa647 | Invitrogen | A31571 | 1:400 |
| Donkey anti-Goat Alexa647 | Invitrogen | A21447 | 1:400 |
| Donkey anti-Goat Alexa488 | Invitrogen | A11055 | 1:400 |

### Supplementary Table 3. Quantitative PCR human primer oligonucleotide sequences.

| Gene name | Forward (5'-3') | Reverse (5'-3') | References |
|---|---|---|---|
| *CFTR* | AGGACTATGGACACTTCGTGCCTT | ATTTGGAACCAGCGCAGTGTTGAC | Ogawa et al. (2021), *Nat Commun* |
| *CK7* | AAGGATGCTCGTGCCAAG | AGCTTCACGCTCATGAGTTC | |
| *SOX9* | TGCATTTCCTCCTGCCTTTGCTTG | GGGCACTTATTGGCTGCTGAAACA | |
| *ALB* | GTGAAACACAAGCCCAAGGCAACA | TCAGCCTTGCAGCACTTCTCTACA | |
| *AFP* | ACAGAGGAACAACTTGAGGCTGTC | AGCAAAGCAGACTTCCTGTTCCTG | |
| *PKD1* | GGACAAGGTGTGAGCCTGAG | AGCTGGTAGACGTCCTCTGT | |
| *PKD2* | TTCCCAGATCAGTCATGGTTTAG | CCTTCCATGCCTTCTGTAGATT | |
| *TMEM16A* | AAGTACTCGACGCTCCCGGCC | ATAAGGAGTTCAGCAGCGTGCCC | |
| *SOX9* | TGCATTTCCTCCTGCCTTTGCTTG | GGGCACTTATTGGCTGCTGAAACA | |
| *SCTR* | TGCATCATGGCCAACTACTC | AATCCCTGGAGGTACTTTCTTTC | |
| *TRPV4* | AGGTGAACTGGTCTCACTGG | GCGAGAAGCCATAATACTGGTAG | |
| *TGR5* | TCTGGCATTGCCCACATT | GAGAAGTTGGGAGCCAAGTAG | |
| *FXR* | ACCACAGATTTCCTCGTCATC | GGCATACGCCTGAGTTCATA | |
| *VANGL1* | TACTCGGACAGAGGAAGTTCAG | GGCAATGTCCTCTTGGGATATG | This paper |
| *VANGL2* | GTCCCAGTACTCGGGCTAT | TCGACTCTTAGAGCGGTGT | |
