## Supplementary Figures for "Bioengineered 3D hPSC-cholangiocyte ducts with physiological signals for biliary disease modelling"

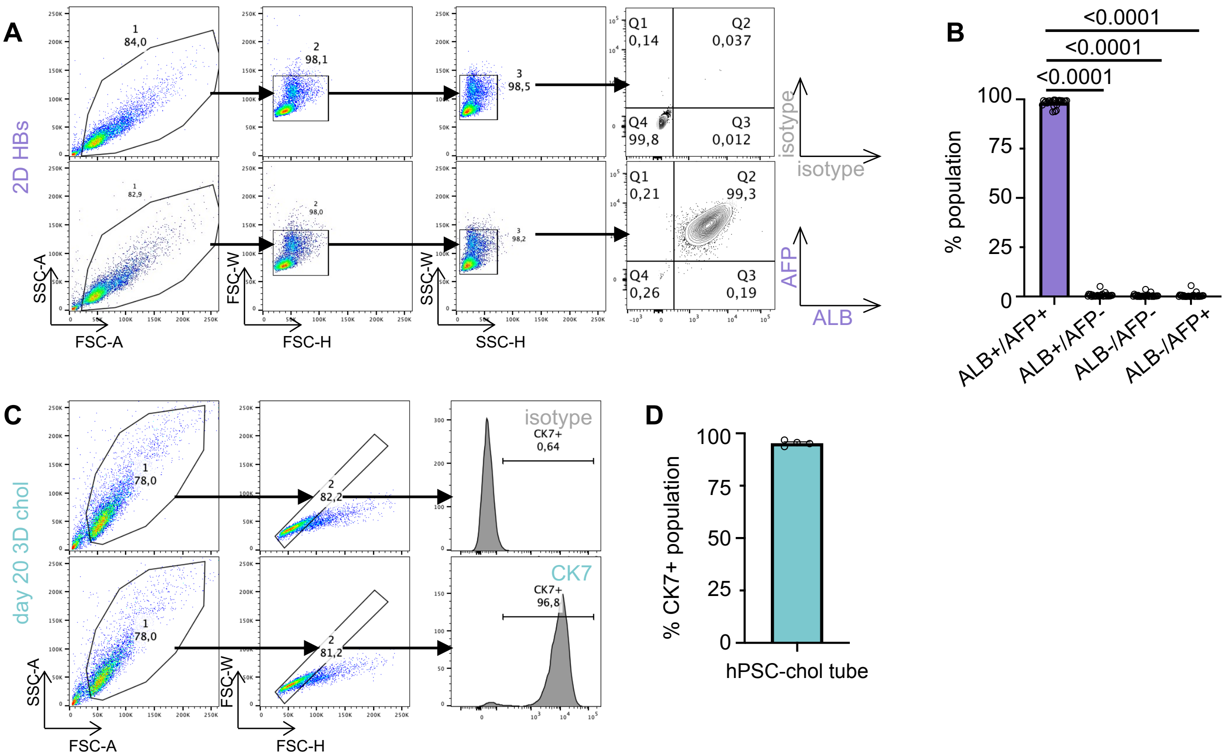


### SUPPLEMENTAL FIGURE 1

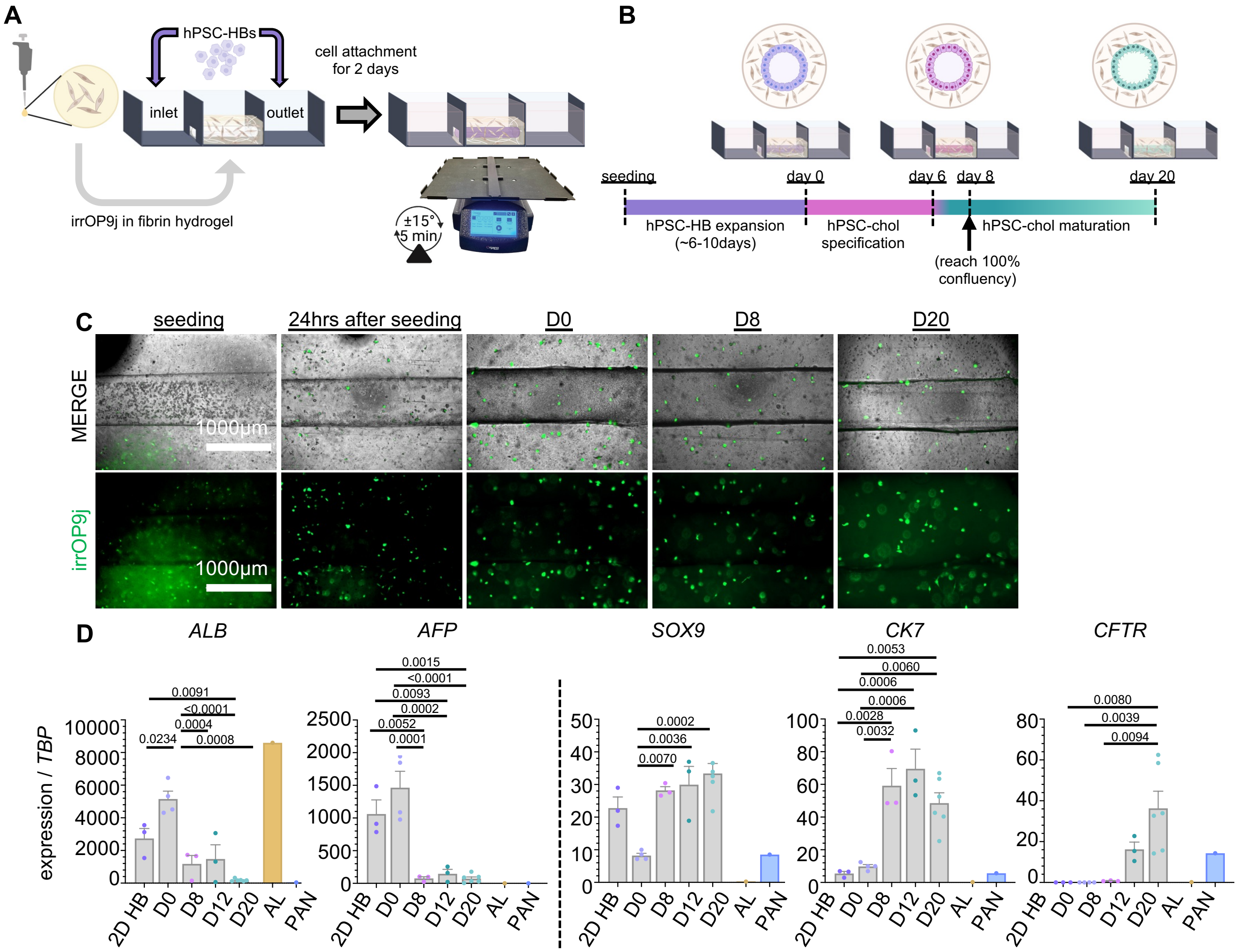


**SUPPLEMENTAL FIGURE 2**

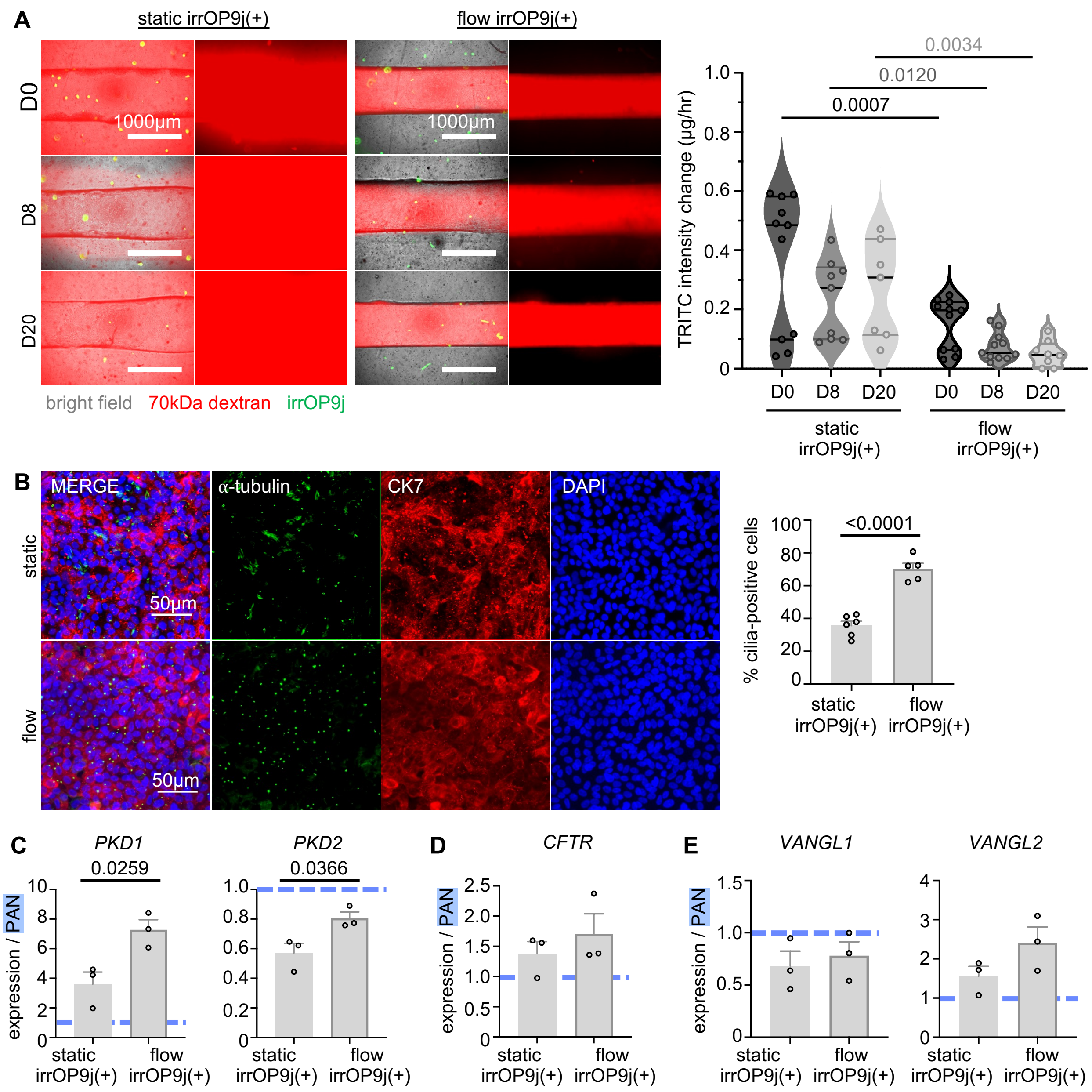


**SUPPLEMENTAL FIGURE 3**

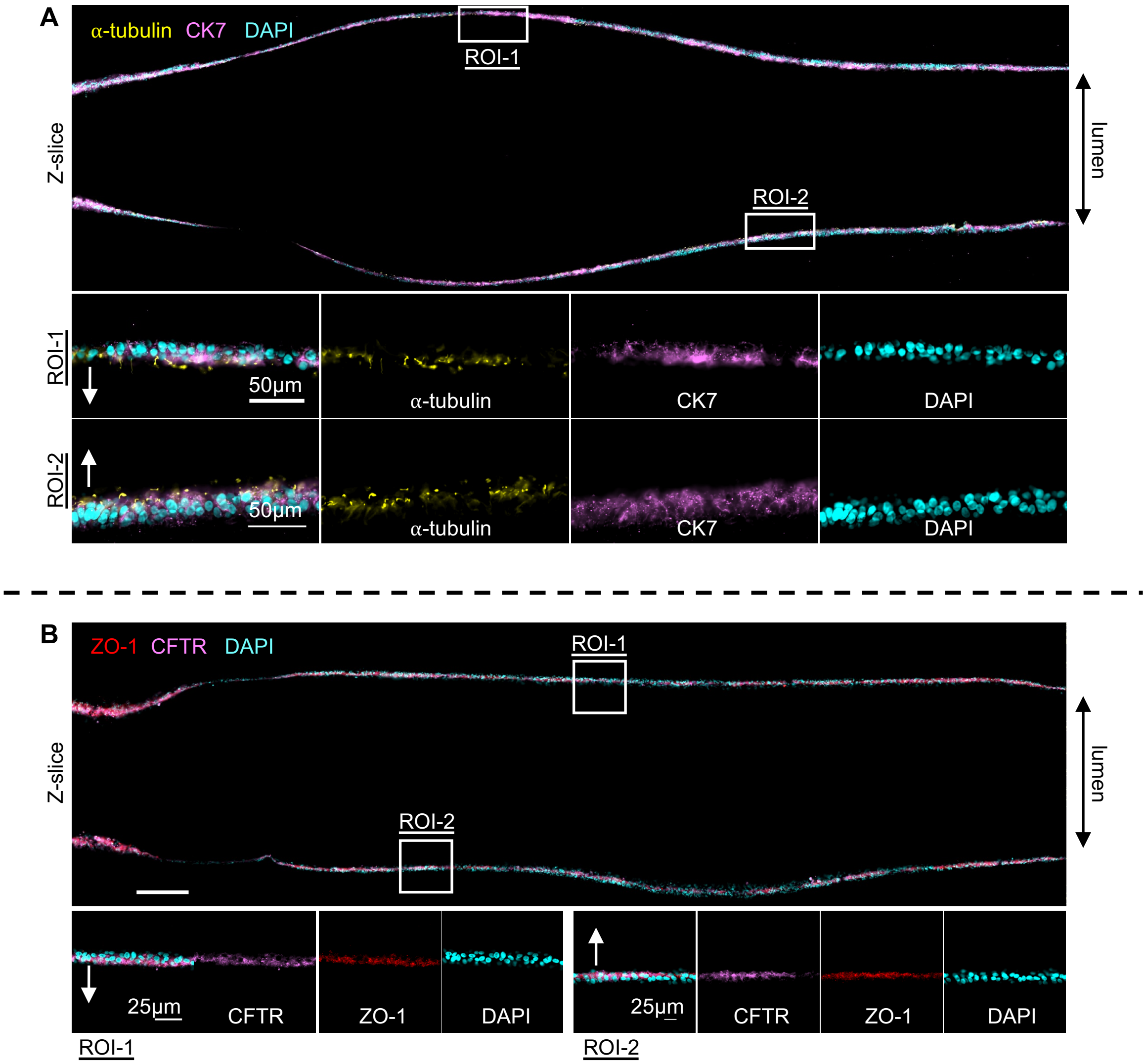


**SUPPLEMENTARY FIGURE 4**

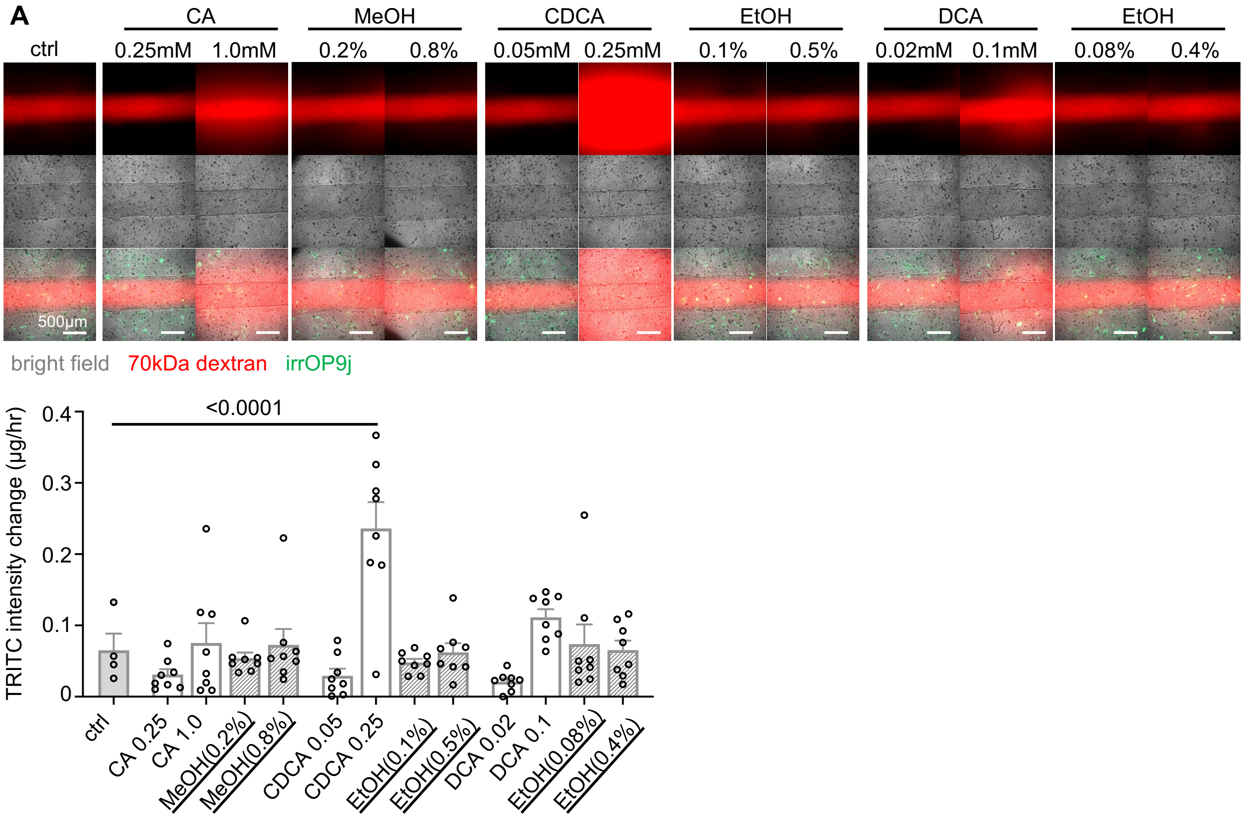


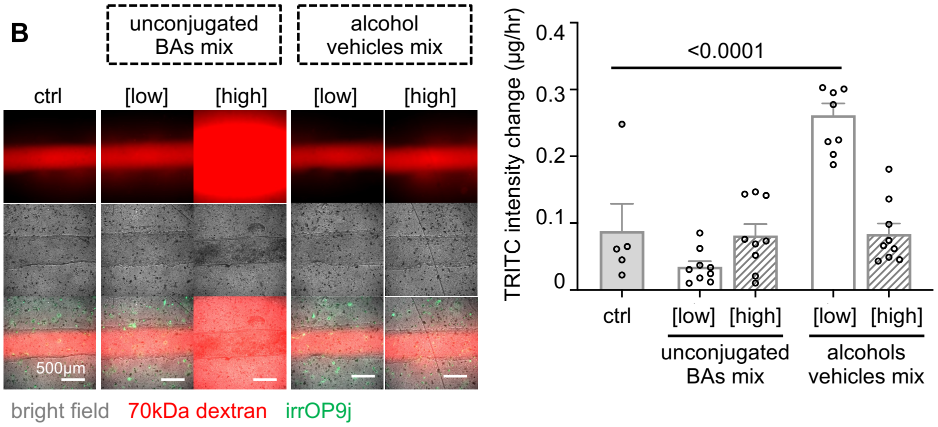


**SUPPLEMENTARY FIGURE 5**

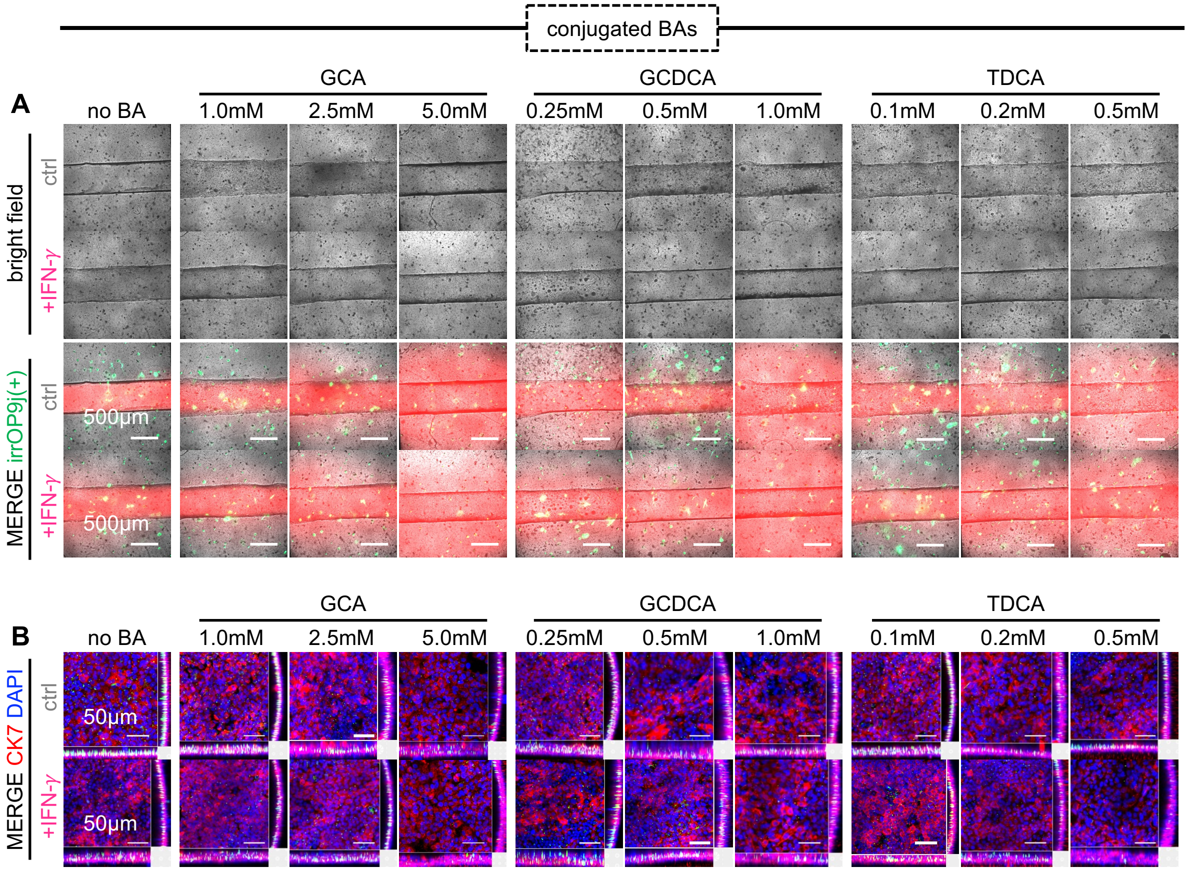


### SUPPLEMENTARY FIGURE 6

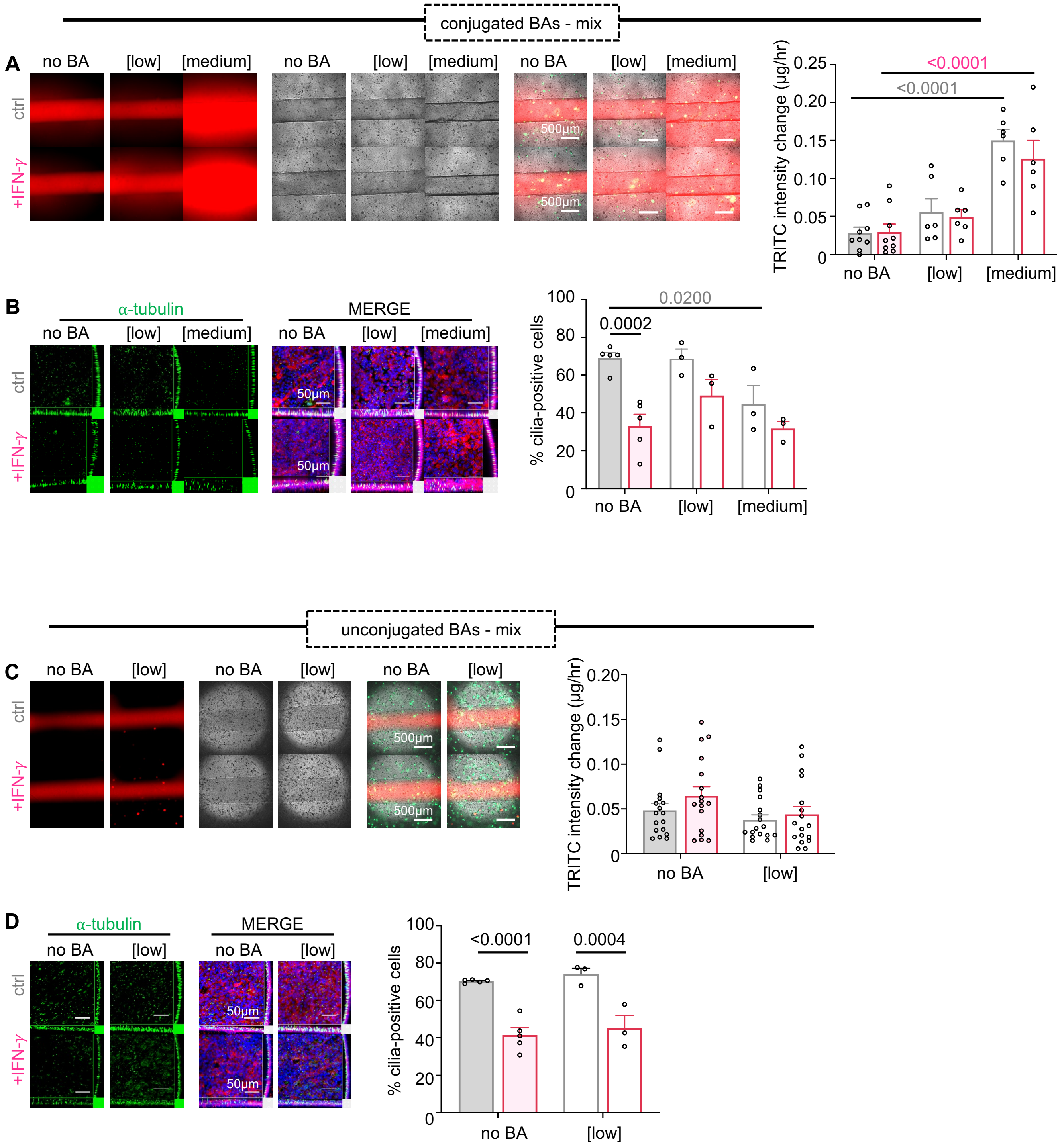


### SUPPLEMENTARY FIGURE 7

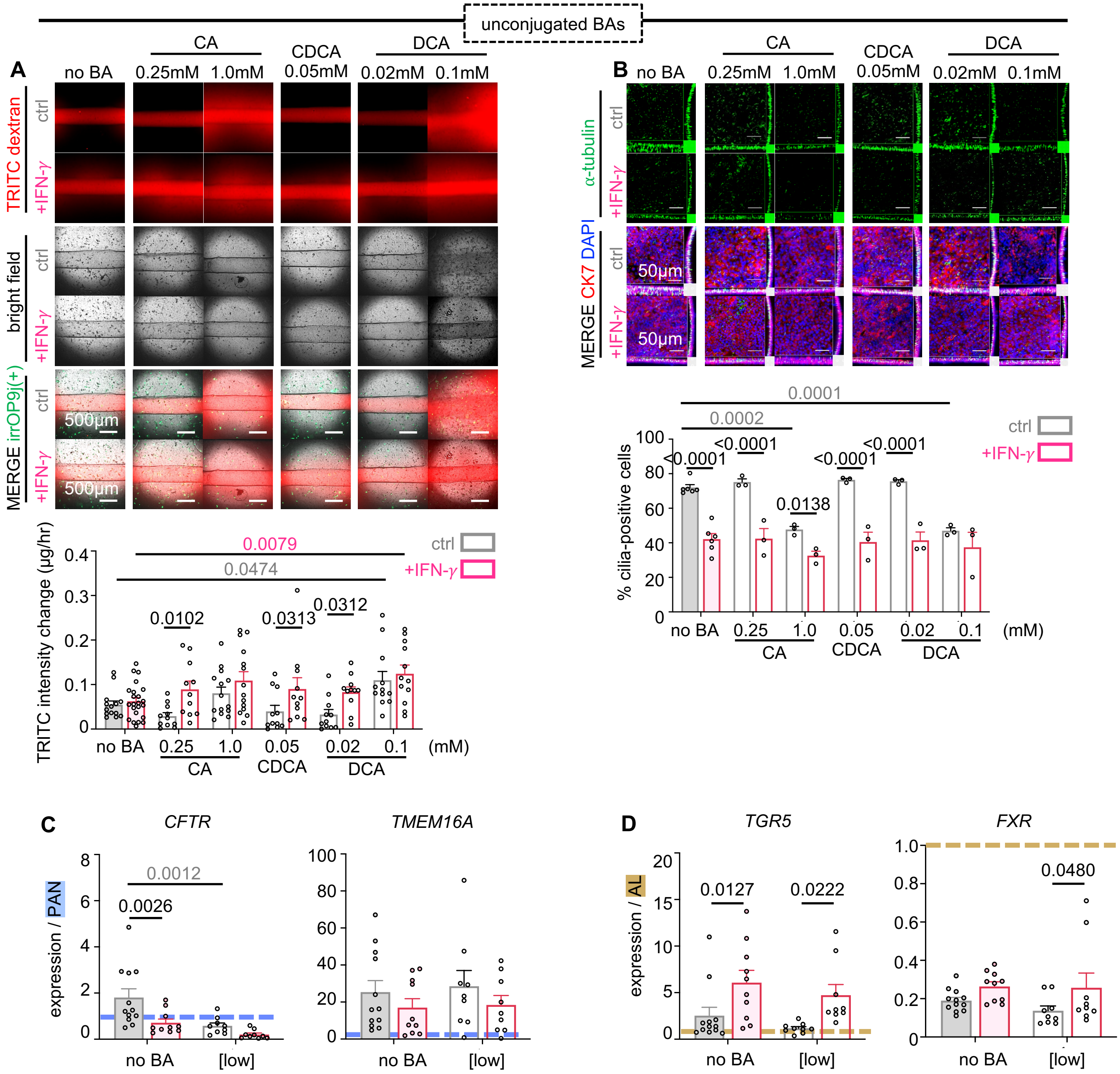


### SUPPLEMENTARY FIGURE 8

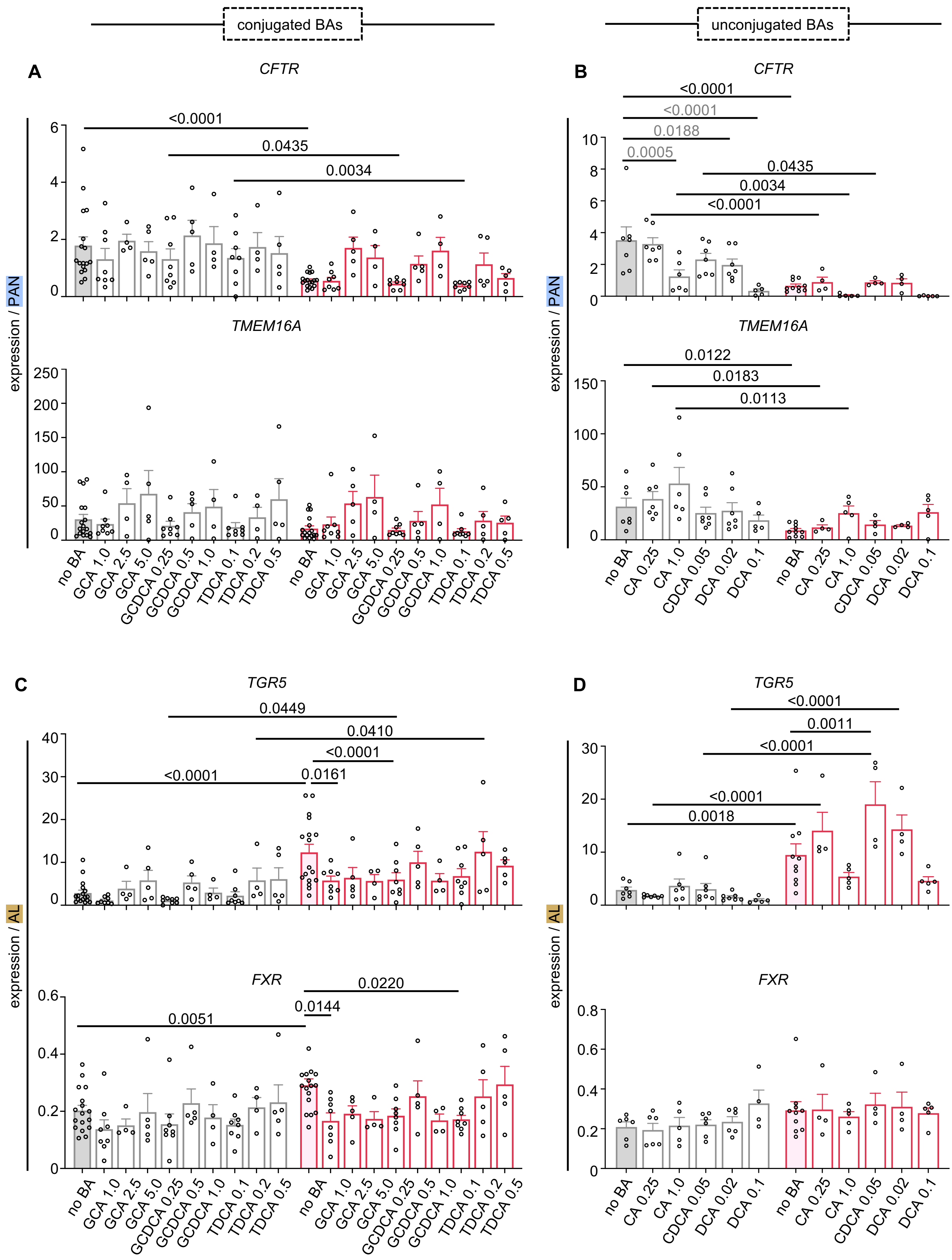


**SUPPLEMENTARY FIGURE 9**

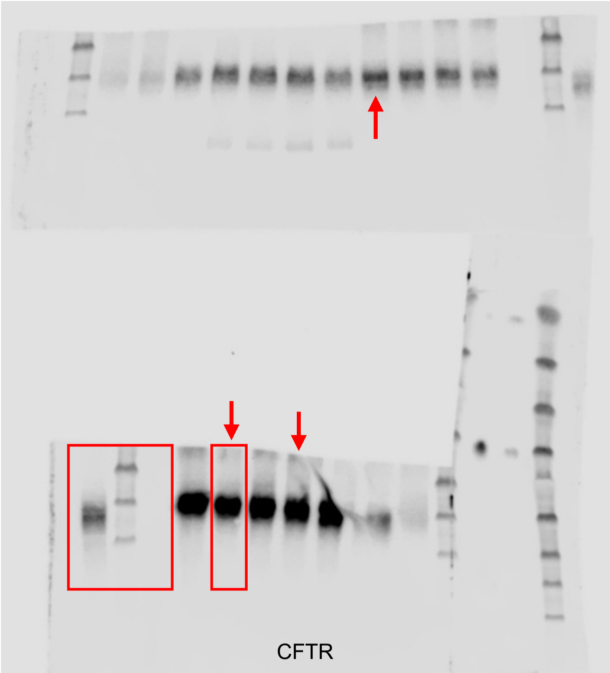


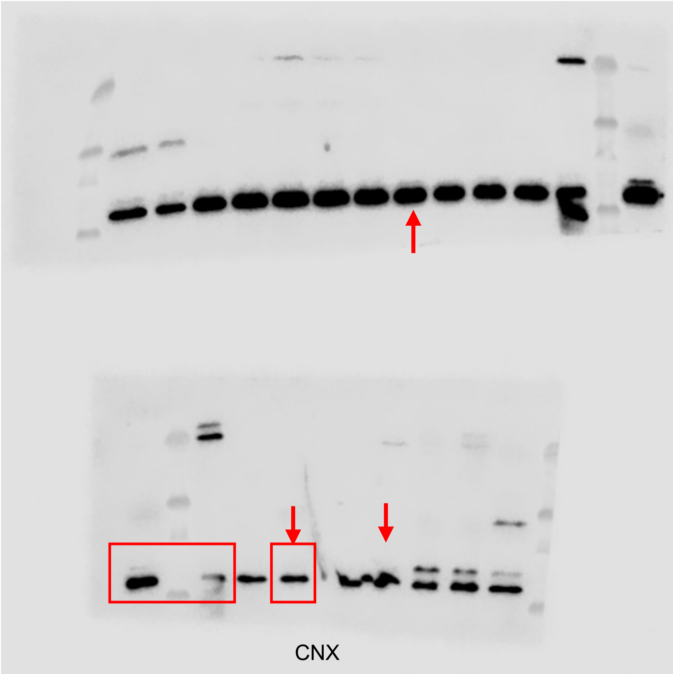


### SUPPLEMENTARY FIGURE 10
